# The shared genetic basis of leaf morphology and tensile resistance underlies the effect of growing season length in a widespread perennial grass

**DOI:** 10.1101/2024.01.10.575100

**Authors:** P. C. Durant, Amit Bhasin, Thomas E. Juenger, Robert W. Heckman

## Abstract

**Premise:** Leaf tensile resistance, a leaf’s ability to withstand pulling forces, is an important determinant of plant ecological strategies. One potential driver of leaf tensile resistance is growing season length. When growing seasons are long, strong leaves—which often require more time and resources to construct than weak leaves—may be more advantageous than when growing seasons are short. Growing season length and other ecological conditions may also impact the morphological traits that underlie leaf tensile resistance.

**Methods:** To understand variation in leaf tensile resistance, we measured size-dependent leaf strength and size-independent leaf toughness in diverse genotypes of the widespread perennial grass *Panicum virgatum* (switchgrass) in a common garden. We then used quantitative genetic approaches to estimate the heritability of leaf tensile resistance and whether there were genetic correlations between leaf tensile resistance and other morphological traits.

**Key Results:** Leaf tensile resistance was positively associated with aboveground biomass (a proxy for fitness). Moreover, both measures of leaf tensile resistance exhibited high heritability and were positively genetically correlated with leaf lamina thickness and leaf mass per area (LMA). Leaf tensile resistance also increased with habitat-of-origin growing season length and this effect was mediated by both LMA and leaf thickness.

**Conclusions:** Differences in growing season length may promote selection for different leaf lifespans and may explain existing variation in leaf tensile resistance in *P. virgatum*. In addition, the high heritability of leaf tensile resistance suggests that *P. virgatum* will be able to respond to climate change as growing seasons lengthen.

## Introduction

Leaf structural resistance is a critical determinant of many ecologically important processes, including leaf longevity, decomposition and turnover, resistance to herbivory, and plant growth rates (Pérez-Harguindeguy et al. 2003). Tough leaves often live longer than weak leaves, which may be especially advantageous when growing seasons are long (Kitajima and Poorter 2010, Kikuzawa et al. 2013). In warm habitats with long growing seasons, leaves have longer to recoup their construction costs, which allows plants to exhibit a slow return on investment strategy (Kudo 1996, Grady et al. 2013, Mason and Donovan 2015). Despite the benefits of high leaf structural resistance, there are also costs—tougher leaves often require more resources, time, and energy to construct than weaker leaves—which can reduce growth rates and thereby decrease performance in resource-rich environments and environments with short growing seasons (Coley et al. 1985, Kitajima and Poorter 2010). Despite its role in many ecological functions, the potential basis of a trade-off between leaf structural resistance and growing season length, including morphological traits and their underlying genetic basis, has been under-explored.

Leaf structural resistance is a key aspect of plant investment in growth and defense, which may depend on growing season length. Long growing seasons can promote greater leaf lifespans in deciduous species (Kikuzawa et al. 2013), potentially leading to the evolution of a conservative functional syndrome—long-lived, low-nutrient leaves with high construction costs, including high leaf mass per area (LMA) (Wright et al. 2004, Onoda et al. 2011, Wang et al. 2023). Similarly, at lower latitudes where growing seasons are typically longer, plants are more likely to experience strong pressure from pathogens and herbivores, which may promote increased allocation to defense (Coley and Barone 1996, Anstett et al. 2016). Plants allocating heavily to defensive traits—including chemical defenses, trichomes, silica content, and structural resistance—often grow slowly and better tolerate abiotic stresses (e.g., Agrawal and Fishbein 2006, Züst and Agrawal 2017, Nardini 2022).

Leaves are exposed to many types of physical damage, including crushing, piercing, pulling, and twisting. Each type of damage exerts a different stress and requires different mechanical properties to resist these stresses. One important mechanical property is leaf tensile resistance—the resistance of a leaf section to being pulled apart—which is analogous to the stress of strong winds or large herbivores pulling on a leaf (Lucas et al. 2000). Measurements of leaf tensile resistance take two forms: size-dependent and size-independent. Size-dependent structural resistance (hereafter, leaf strength) tends to increase with leaf size (width, length, and area) for any given leaf composition—it takes more energy to break through more leaf material (Lucas et al. 2000). Size-independent structural resistance (hereafter, leaf toughness), on the other hand, is a material property related primarily to leaf composition, especially cell wall composition, leaf thickness, or leaf density (Westbrook et al. 2011). Some commonly studied leaf functional traits, like leaf mass per area (LMA) and leaf thickness, have also been shown to contribute to leaf tensile resistance in broad multi-species surveys (Onoda et al. 2011).

To date, most studies investigating the basis of leaf structural resistance have examined patterns across many species (Westbrook et al. 2011, Onoda et al. 2017). While this cross-species approach is valuable, it has limited ability to identify the drivers of current selection on leaf structural resistance, which is best accomplished within species (Agrawal 2020). Moreover, working within a single species allows researchers to use the powerful suite of quantitative genetics tools. For instance, quantitative genetics methods can be used to examine the capacity of a trait like leaf structural resistance to evolve predictably within a population or respond to artificial selection imposed by breeding programs. Predictable evolution requires two things: natural selection acting on the trait and standing genetic variation in the trait (Lynch and Walsh 1998, Mackay 2001). Traits exhibiting little genetic variation within the population (i.e., low heritability) and/or traits under limited natural selection will evolve more slowly than traits with a high heritability under strong natural selection (Lynch and Walsh 1998). Quantitative genetics can also be used to examine the anatomical or compositional traits that best explain leaf structural resistance. Strong genetic correlations can result from genetic linkage (nearby loci are frequently inherited together), pleiotropy (a locus that affects multiple traits), or population structure (non-random mating within a population; Lynch and Walsh 1998, Walsh and Blows 2009). While heritability and genetic correlation describe the aggregate genetic contribution to traits or pairs of traits, genome-wide association (GWA) mapping can identify particular genomic regions associated with leaf structural resistance and whether any genomic regions are associated with both leaf structural resistance and leaf composition.

In order to study the genetic mechanisms underlying leaf structural resistance, it is advantageous to examine a species with a broad geographic distribution, large morphological differences, and adequate genetic resources. One such species is *Panicum virgatum* (switchgrass), a perennial grass whose distribution encompasses much of eastern and central North America, where it experiences considerable variation in growing season length, potentially leading to differing selective pressures on leaf strength (Casler 2012, Lovell et al. 2021). We take advantage of this broad geographic extent to examine the geographic, anatomical, and genetic drivers of leaf tensile resistance in a diverse group of resequenced *P. virgatum* genotypes growing in a common location. We first assessed the links between growing season length and leaf structural resistance, then examined whether genetic or phenotypic relationships existed between leaf structural resistance traits and leaf traits that characterize the size or construction of the leaf (LMA, leaf lamina thickness, and leaf density). This allowed us to assess four questions: 1) How much phenotypic and genetic variation in leaf tensile resistance is present within *P. virgatum*? 2) Is there evidence for selection for leaf tensile resistance in *P. virgatum*? 3) Which leaf morphological traits contribute to leaf tensile resistance and do these traits share a genetic basis? 4) How do leaf morphology and its contribution to leaf tensile resistance change in response to growing season length?

## Methods

### Study system, sequencing, and experimental design

*Panicum virgatum* is a morphologically diverse, widespread C_4_ grass species whose range spans much of eastern and central North America, from southern Canada to Mexico (Casler et al. 2011, Lovell et al. 2021). The species has three genetically and geographically distinct subpopulations—Gulf, Midwest, and Atlantic (Lovell et al. 2021). The Gulf subpopulation occupies the southernmost portion of the US range; it is adapted to long growing seasons and has large, long-lived leaves (Lovell et al. 2021). The Midwest subpopulation occurs in the north-central United States in upland environments; it is adapted to short growing seasons and has small, short-lived leaves (Lovell et al. 2021). The Atlantic subpopulation is found along the eastern seaboard; it has leaf attributes that are intermediate between the other subpopulations (Lovell et al. 2021).

The *P. virgatum* genotypes sampled in this study came from natural occurrences and cultivars collected from across the United States between 2010 and 2018 (Fig. 1). The genotypes represented all three genetic subpopulations identified by Lovell et al. (2021) and their collection sites covered 18.5 degrees of latitude. Prior to planting, all genotypes were propagated at the Brackenridge Field Laboratory, University of Texas at Austin (Austin, TX, USA). Each genotype was grown in an 18.9-L pot until the crown was large enough to be divided into multiple plants, each of which was then grown in a 3.78-L pot. One clone of each genotype was then transported to the J.J. Pickle Research Campus at the University of Texas at Austin (Austin, TX, USA) in May 2018, where it was planted in a common garden covered in landscape fabric (Sunbelt 3.2 oz, DeWitt, Sikeston, MO, USA). Genotypes in the common garden were randomly arranged 1.56 m apart in a honeycomb pattern. The edges of the common garden were surrounded by a row of plants of Blackwell, a common upland *P. virgatum* cultivar. Of the 456 genotypes we surveyed, 165 were from the Atlantic subpopulation, 201 were from the Gulf subpopulation, and 90 were from the Midwest subpopulation.

**Fig. 1.**
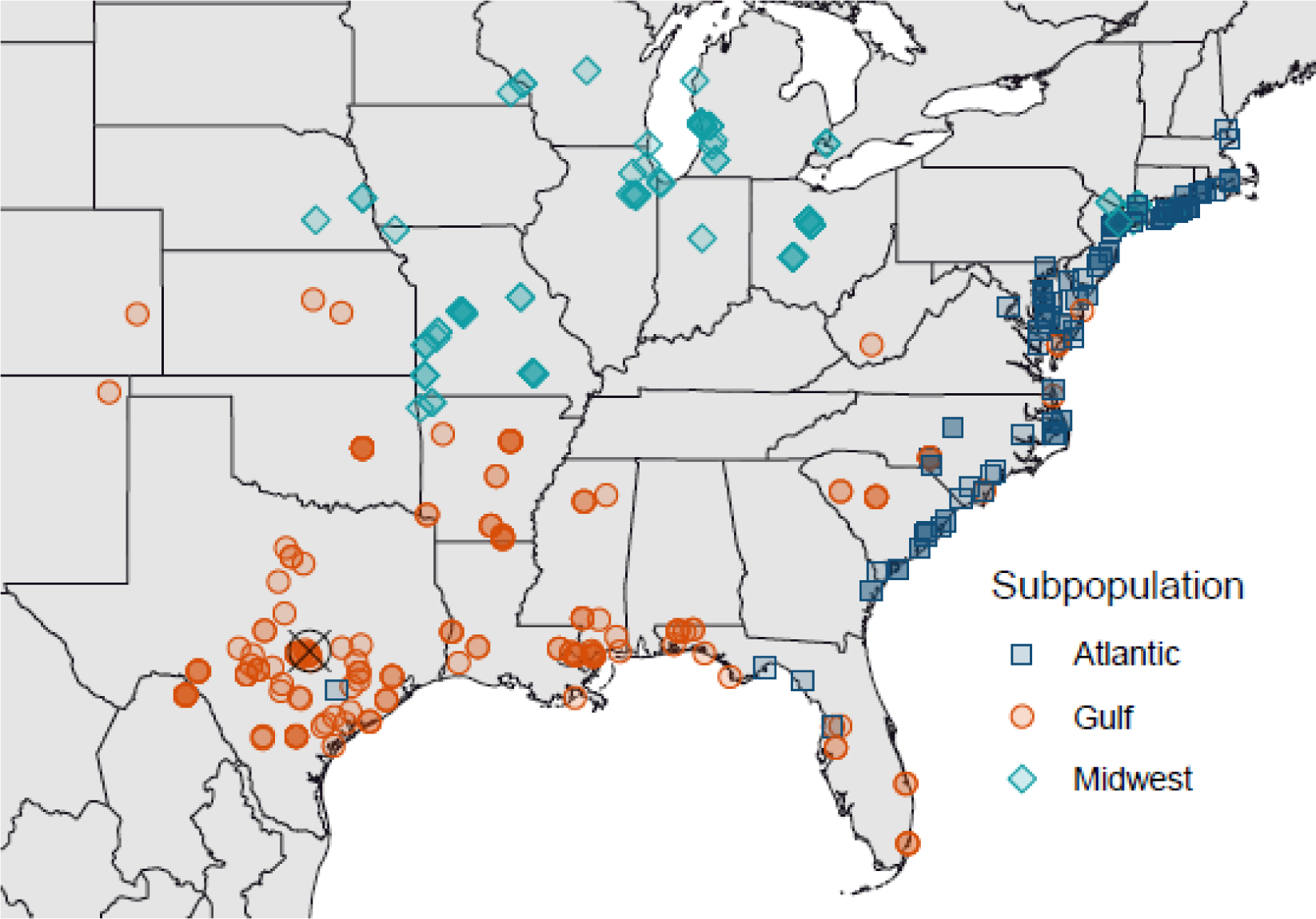
Locations of origin for genotypes used in this study. All genotypes were propagated at the University of Texas at Austin, then planted in a common garden in Austin, TX, USA (denoted by a circle cross). *Panicum virgatum* has three genetic subpopulations, which are largely geographically separated.

The methods for sequencing and annotating the switchgrass genome and determining the genetic subpopulation for each genotype are discussed in detail by Lovell et al. (2021). Briefly, the sequenced *P. virgatum* genome came from a genotype of the Alamo cultivar (AP13; version 5.1, https://phytozome-next.jgi.doe.gov/info/Pvirgatum_v5_1). All genotypes were resequenced (Illumina HiSeq X10 and Illumina NovaSeq 6000 paired-end sequencing, 2 × 150 bp) and mapped against the reference genome (bwa-mem; Li and Durbin 2009). 10.2 million single nucleotide polymorphisms (SNPs) remained after removing missing data and loci with minor allele frequencies < 0.05. These SNPs were used in the analyses described below.

### Phenotypic data collection

To evaluate leaf structural resistance in *P. virgatum*, we measured four leaf attributes— leaf strength, leaf width, leaf mass per area (LMA), and leaf thickness—which we then used to estimate two additional attributes—leaf toughness and leaf density. To accomplish this, we collected three young, fully expanded leaves from each plant that were representative of the entire canopy. Leaves were rehydrated by placing their cut end in 15-mL tubes filled with ∼ 5 mL of water, then stored in cool, dark conditions for up to three days. Maintaining consistent hydration across samples can be important because the tensile strength of plant tissue can change with moisture content (e.g., Hales and Miniat 2017).

Leaf strength was the maximum tensile force that a leaf could withstand before failing (a size-dependent quantity). The leaf strength assay involved conducting a constant displacement rate tensile strength test on rectangular leaf segments with a tensiometer (Instron Model 1000 Universal Testing Machine, Instron, Norwood, MA). Specifically, we first cut a 20-cm-long segment of each leaf; we avoided the ends of the leaf, where leaf width tapers rapidly. Because the midrib of *P. virgatum* is considerably stronger than the leaf lamina, we weakened the midrib with a small incision ∼5 cm away from the basal end of the leaf segment. To protect the tips of leaf segments from the tensiometer clamps, we covered them with ethylene-vinyl acetate craft foam. The arm of the tensiometer moved at a constant displacement rate of 0.01 mm s^-1^ until the leaf segment completely failed in tension. During the test, the tensile load and total displacement were collected using a 250 N load cell and a digital displacement sensor that is built-in at the top of the actuator. When a leaf experienced multiple fractures (e.g., when the leaf initially tore only partially), we still considered the maximum force experienced by the sample as the measure of leaf strength. Leaf toughness was the ratio of the maximum load (leaf strength) to leaf width at the fracture point, which was 5 cm from the basal end of the leaf segment 72.7% of the time. Thus, leaf toughness normalizes the influence of varying leaf width, but would still be influenced by leaf thickness and intrinsic strength.

We also measured two leaf morphological traits that are frequently associated with leaf tensile resistance: leaf mass per area (LMA) and leaf thickness. LMA is the ratio of dried leaf mass to fresh, one-sided leaf area. Leaf area was estimated using a leaf area meter (LI-3100C, LI-COR Biosciences, Lincoln, NE). Leaves were then dried at 50 °C for 72 h and weighed. On these leaves, we also recorded the thickness of the leaf lamina at 5 cm from the leaf base and at the approximate midpoint along the leaf length. The average of the thickness near the base and at the middle of the leaves was used in analyses. From these measurements, we then estimated leaf density: the ratio of LMA to the average of leaf lamina thickness (base and midpoint). Because leaf strength and LMA measurements are both destructive, we measured leaf strength on one leaf and LMA and leaf thickness on two other leaves from the same plant.

In 2019, 2020, and 2021, plants were harvested each fall at ∼ 15 cm above ground level and aboveground biomass was weighed to estimate biomass production. A representative subsample of vegetation from each plant was collected and weighed fresh, then dried and reweighed; the percent moisture in this subsample was used to calculate the dry mass of the total plant. We then summed biomass for each plant across the three years. Biomass measured in 2019 was also reported by Lovell et al. (2021).

### Analyses

We performed all analyses using R 4.2.2 (R Core Team, 2021). To address our first question—how much phenotypic and genetic variation in leaf tensile resistance is present within *P. virgatum*—we took two approaches. First, we examined phenotypic differences in leaf strength and leaf toughness among genetic subpopulations using ANOVA and post-hoc Tukey’s Honestly Significant Difference tests (nlme and emmeans packages; Pinheiro and Bates 2000, Lenth 2023, Pinheiro et al. 2023). Second, we examined the genetic contributions to traits using a linear mixed model in which the random effect is an additive genetic relationship matrix (animal model, ASReml-R package; Wilson et al. 2010, Butler 2021). The additive genetic relationship matrix was calculated using the vanRaden method—the similarity between each pair of plants was calculated at each locus (i.e., SNP) and weighted by the minor allele frequency of the locus, then summed across all loci (VanRaden 2008, Lovell et al. 2021). We calculated narrow sense heritability (*h*^2^) in univariate-response models as the ratio of additive genetic variance (V_A_) to total variance (V_A_ + residual variance, V_ɛ_). This analysis was performed for all genotypes together as well as separately within each genetic subpopulation. Prior to analysis, we mean-centered and variance-standardized (mean = 0, standard deviation = 1) responses.

To address our second question—is there evidence for selection for leaf tensile resistance in *P. virgatum*—we performed a selection gradient analysis (Lande and Arnold 1983). To do this, we first mean-standardized aboveground biomass for each genotype by dividing each genotype’s biomass by the global mean aboveground biomass. Previous work demonstrates that aboveground biomass is highly correlated with seed production in *P. virgatum* (Palik et al. 2016), making it a useful proxy for fitness (Lowry et al. 2019). This results in a relativized score where average fitness is equal to 1, above average fitness > 1, and below average fitness < 1 (Franklin and Morrissey 2017). We then examined how mean-centered and variance-standardized (mean = 0, standard deviation = 1) leaf toughness or leaf strength predicted mean-standardized biomass in a single generalized least squares model (gls() function in package nlme; Pinheiro and Bates 2000, Pinheiro et al. 2023). The selection gradient model included linear and quadratic effects of leaf strength or leaf toughness and their interaction with subpopulation. All values were standardized separately by subpopulation. Quadratic effects were doubled prior to analysis (Stinchcombe et al. 2008).

To address our third question—which leaf morphological traits contribute to leaf tensile resistance, and do these traits share a genetic basis—we took two approaches. First, we calculated the phenotypic correlations (Pearson’s correlation coefficient, r_P_; cor() function) and genetic correlations (r_G_) between leaf strength or leaf toughness and leaf thickness, LMA, and leaf density in bivariate-response linear mixed models with the additive genetic relationship matrix as a random effect. r_G_ was the ratio of the genetic covariance (cov_1,2_) between a pair of traits to the square root of the product of the additive genetic variances of the two traits—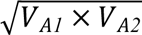 (Wilson et al. 2010). We quantified the significance of r_G_ using a likelihood ratio test to compare a model that freely estimated the genetic covariance between traits to a model that fixed their genetic covariance at 0. As with the univariate mixed model analyses described above, we analyzed all genotypes together, and we mean-centered and variance-standardized all responses.

Then, we performed association mapping (GWA) for each leaf trait (leaf area, leaf thickness, LMA, leaf density, leaf strength, and leaf toughness) with linear models that included 10.2 million biallelic SNPs (switchgrassGWAS and bigsnpr packages; Privé et al. 2018, Lovell et al. 2021). In each model, we accounted for population structure (Q, sensu Yu et al. 2006) by including one to 15 principal components of the SNP matrix and selecting the number of principal components that resulted in the best model fit (λ_GC_ nearest 1 calculated from quantile-quantile plots). We used a 5% false discovery rate significance threshold (Storey and Tibshirani 2003) to identify significant associations between leaf traits and SNPs. Quantile-quantile plots of expected and observed p-value distributions for each trait are available in the supplementary material (Fig. S1).

To identify significant associations between groups of leaf traits and SNPs, we used a data fusion method called the Cauchy combination test (CCT), which is a computationally efficient method with no assumption of independence between responses (ACAT package; Liu 2020). Specifically, we examined associations between each SNP and 1) both tensile traits (leaf strength and leaf toughness), 2) pairs of leaf tensile and leaf morphology traits, and 3) all five leaf traits. A significant association would provide evidence that pleiotropic changes in leaf morphology can impact leaf tensile resistance. Despite the computational efficiency of the CCT, we still needed to reduce the number of SNPs tested to alleviate the computational burden (Kremling et al. 2019, Wu et al. 2022). To do this, we first identified the minimum p-value for each SNP across all five traits in univariate GWAS, then selected the 10% of SNPs with the lowest minimum p-value. Because we performed only a portion of the total possible tests, we could not calculate the 5% false discovery rate significance threshold. Instead, we used the more stringent Bonferroni correction (ɑ = 0.05, *m* equal to 10.2 million biallelic SNPs) to reduce Type I errors.

To identify candidate genes associated with leaf traits for both CCT and single-trait GWA, we investigated the genes within 10 kbp of each SNP and their orthologous functions in the model plants *Arabidopsis thaliana* and *Oryza sativa*.

Finally, to address our fourth question—how do leaf morphology and its contribution to leaf tensile resistance change in response to growing season length—we performed mediation analysis, which is a method for assessing the importance of indirect effects on a response (Zhao et al. 2010). Specifically, we examined whether the effect of growing season length on leaf tensile resistance was mediated by the effects of leaf morphology in a multivariate model (i.e., structural equation model, lavaan package; Rosseel 2012). We estimated growing season length by calculating growing degree days for the site-of-origin of each genotype—the sum of each day’s mean temperature minus 12 °C, when mean daily temperature exceeds 12 °C. Mean daily temperature was the average of daily maximum and minimum temperature at each site-of-origin for 2001–2019 (package nasapower; Sparks 2018, 2023). This model had two levels—the effect of growing degree days on LMA and leaf thickness and the effects of LMA and leaf thickness on leaf toughness or leaf strength (sem() function in package lavaan). Following Zhao et al. (2010), we estimated the indirect effect of growing degree days on leaf strength or leaf toughness by multiplying two paths—the effect of growing degree days on LMA or leaf thickness and the effect of LMA or leaf thickness on leaf strength or leaf toughness—then estimated the significance of these effects with 10,000 bootstrapping iterations. A significant effect indicates that the relationship between growing season length and leaf tensile resistance is mediated by leaf morphology (Zhao et al. 2010). We then examined whether this mediation differs among genetic subpopulations using a multigroup model. Both the single group and multigroup models also included a correlation between LMA and leaf thickness.

## Results

### Leaf tensile resistance is variable and highly heritable in switchgrass

Leaf strength and leaf toughness both differed significantly among genetic subpopulations (p < 0.001; Table 1, Fig. 2A, B). Specifically, Gulf subpopulation leaves were more than twice as strong as Atlantic and Midwest leaves (109% and 198% stronger, respectively; p < 0.001). Additionally, Atlantic leaves were 43% stronger, on average, than Midwest leaves (p < 0.001). As with leaf strength, Gulf subpopulation leaves were significantly tougher than Atlantic and Midwest leaves; however, the differences in leaf toughness were of considerably smaller magnitude—Gulf leaves were 37% tougher than Atlantic leaves (p < 0.001) and 36% tougher than Midwest leaves (p < 0.001). The toughness of Atlantic and Midwest leaves did not differ (p = 0.97; Table 1, Fig. 2B). These smaller subpopulation differences in leaf toughness were likely driven by leaf size—Gulf leaves were 71% larger than Atlantic leaves and 189% larger than Midwest leaves (Table 1)—rather than leaf composition. Both leaf strength and leaf toughness were highly heritable across all genotypes and in the Gulf and Atlantic subpopulations (h^2^ > 0.5), but less heritable in the Midwest subpopulation (h^2^ < 0.3; Table 1).

**Fig. 2.**
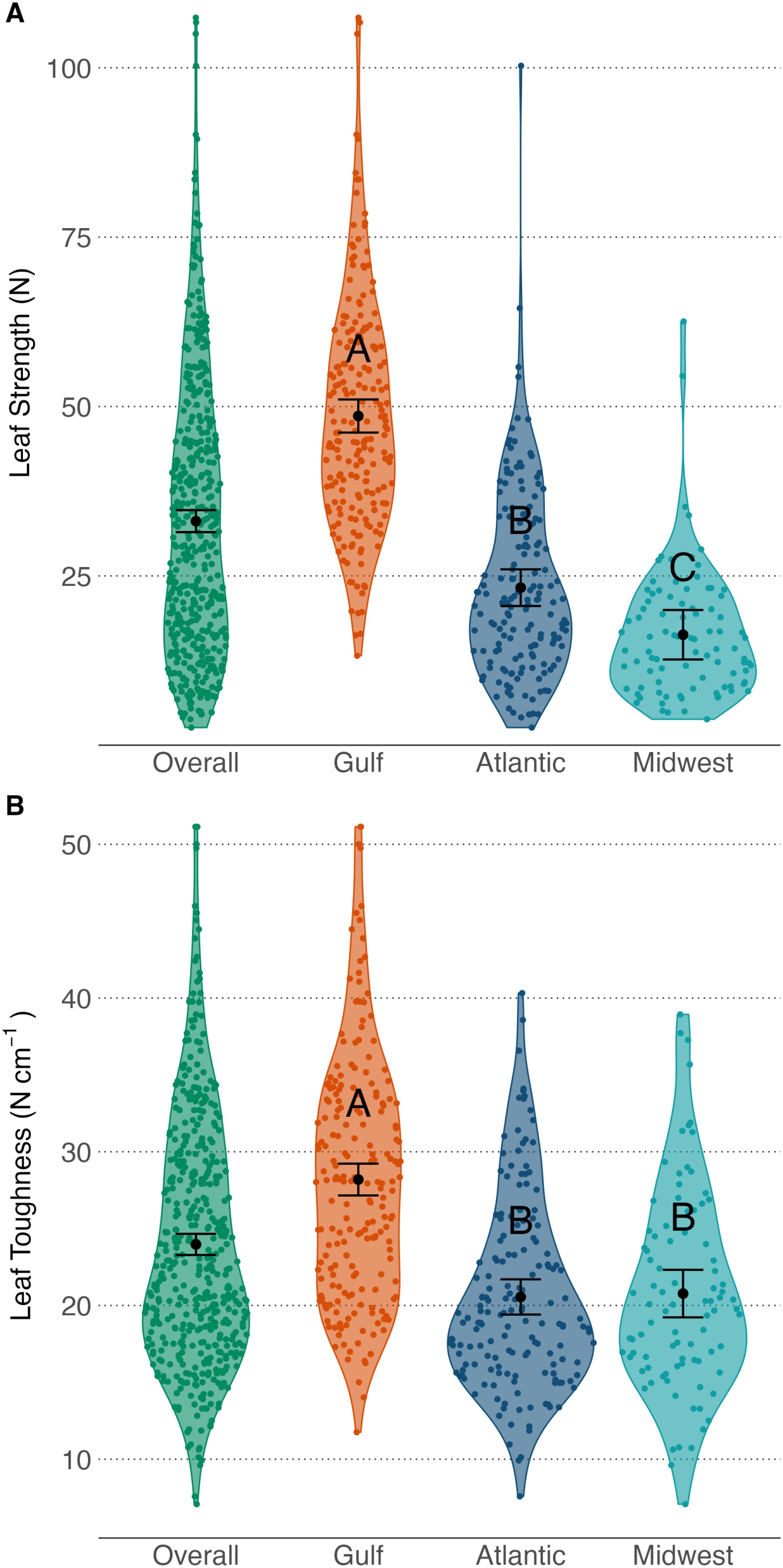
Differences in A) leaf strength and B) leaf toughness among populations of *P. virgatum*. Means and 95% confidence intervals, shown by error bars, were model-derived. Points are raw values that were jittered horizontally to reduce overplotting.

**Fig. 3.**
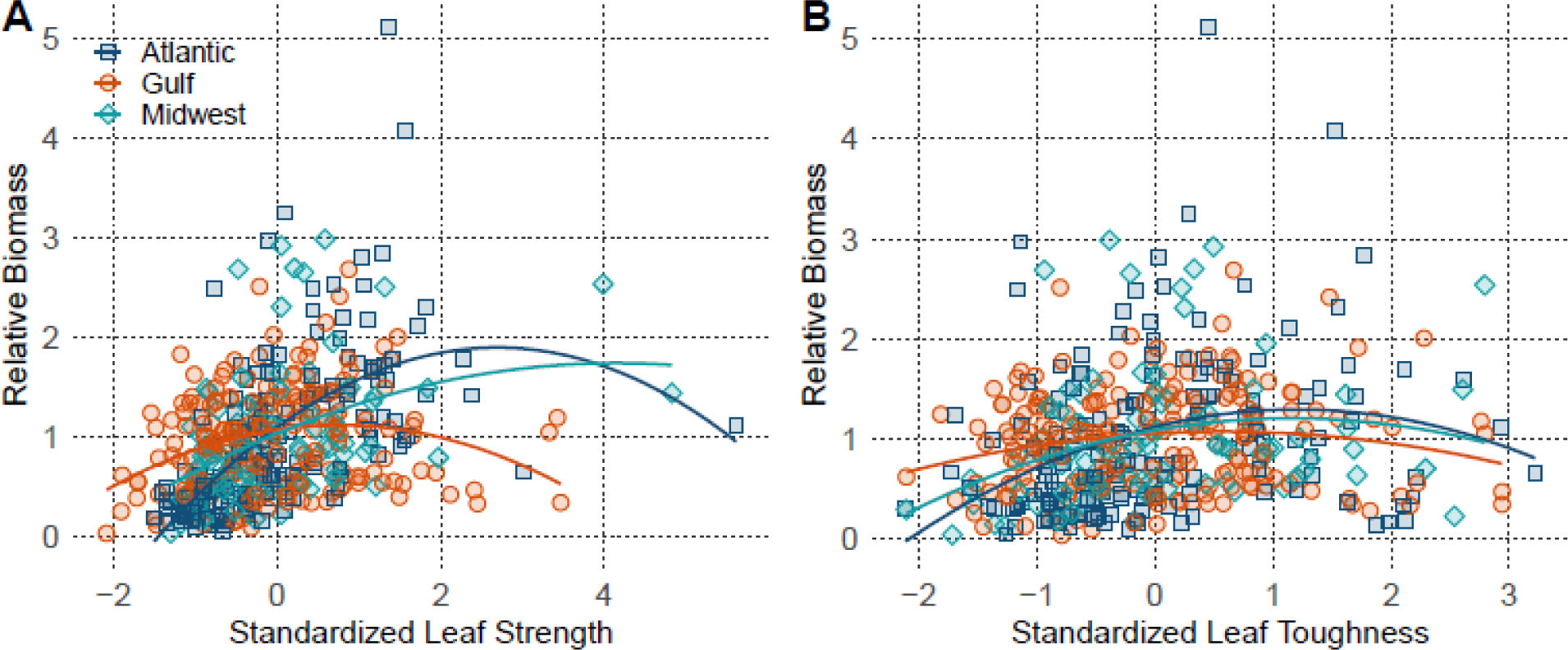
Effects of A) standardized leaf strength and B) standardized leaf toughness (transformed such that across all plants within a subpopulation, mean = 0 and standard deviation = 1) on relative biomass. Model-derived quadratic parameter estimates are doubled. Colors and shapes are used to differentiate subpopulations.

**Table 1.** Phenotypic mean and narrow-sense heritability of leaf traits. Heritability (*h^2^*) was calculated from a univariate model with an additive genetic relationship matrix as a random effect. Standard errors (SE) were calculated using the delta method.

| | | Mean ± SE | Narrow-sense<br>heritability ($h^2$) ±<br>SE |
| --- | --- | --- | --- |
| Strength (N) | Overall | 33.1 ± 0.8 | 0.87 ± 0.04 |
| | Gulf | 48.6 ± 1.2 | 0.89 ± 0.05 |
| | Atlantic | 23.3 ± 1.4 | 0.87 ± 0.09 |
| | Midwest | 16.3 ± 1.9 | 0.19 ± 0.30 |
| Toughness (N cm<sup>-1</sup>) | Overall | 24.0 ± 0.4 | 0.76 ± 0.08 |
| | Gulf | 28.2 ± 0.5 | 0.83 ± 0.08 |
| | Atlantic | 20.6 ± 0.6 | 0.54 ± 0.03 |
| | Midwest | 20.8 ± 0.8 | 0.27 ± 0.35 |
| Leaf thickness (mm) | Overall | 0.224 ± 0.002 | 0.75 ± 0.08 |
| | Gulf | 0.265 ± 0.003 | 0.85 ± 0.07 |
| | Atlantic | 0.197 ± 0.004 | 0.64 ± 0.20 |
| | Midwest | 0.185 ± 0.005 | 0.07 ± 0.18 |
| LMA (g m<sup>-2</sup>) | Overall | 875 ± 9 | 0.81 ± 0.08 |
| | Gulf | 1009 ± 11 | -- |
| | Atlantic | 782 ± 11 | -- |
| | Midwest | 742 ± 12 | -- |
| Leaf density (g cm<sup>-3</sup>) | Overall | 0.398 ± 0.003 | 0.53 ± 0.15 |
| | Gulf | 0.388 ± 0.005 | 0.60 ± 0.18 |
| | Atlantic | 0.404 ± 0.005 | 0.25 ± 0.26 |
| | Midwest | 0.407 ± 0.006 | 0.34 ± 0.25 |

### Leaf tensile resistance is associated with biomass within and across subpopulations

In general, leaf tensile resistance was positively associated with plant biomass, a proxy for fitness (Table S1, S2; Fig. S2, 3). Specifically, the linear effect of leaf strength on plant biomass was significantly greater than 0 in all three subpopulations (p < 0.01 for each subpopulation; Table S3A). This effect was strongest in the Atlantic subpopulation (Atlantic – Gulf, p < 0.001; Atlantic – Midwest, p = 0.032) and marginally stronger in the Midwest subpopulation than in the Gulf subpopulation (p = 0.056; Table S3A). There was also an overall significant quadratic effect of leaf strength on biomass that was concave down (p < 0.001; Table S1A, S2A), indicating that the overall relationship between biomass and leaf toughness is non-linear and saturating. Within subpopulations, this quadratic effect was significant in the Atlantic and Gulf subpopulations (p < 0.001 and p = 0.003, respectively), but not in the Midwest subpopulation (p = 0.15; Table S3A); none of the subpopulations differed significantly from one another (Table S4A).

Like leaf strength, leaf toughness also had a positive association with plant biomass across all populations (p < 0.001) and the effect differed among subpopulations (p = 0.029; Table S1B). Specifically, biomass increased with leaf toughness in the Atlantic (p < 0.001) and Midwest (p = 0.006), but not in the Gulf subpopulation (p = 0.201; Table S3B). This relationship was significantly stronger in the Atlantic than Gulf subpopulation (p = 0.012), but did not differ between other subpopulations (Atlantic – Midwest, p = 0.67; Gulf – Midwest, p = 0.26; Table S4B). As with leaf strength, there was also an overall quadratic effect of leaf toughness on biomass that was concave down (p < 0.001; Table S1B, S2B), indicating a saturating effect of leaf toughness on biomass. Within subpopulations, this quadratic effect was significant in the Atlantic subpopulation (p = 0.002), but not in the Gulf or Midwest subpopulations (p = 0.101 and p = 0.076, respectively; Table S3B); none of the subpopulations differed significantly from one another (Table S4B). These results indicate that leaf tensile resistance can be an important determinant of plant fitness, but that selection on leaf tensile resistance can differ considerably among genetic subpopulations.

### Leaf size and LMA are correlated with leaf tensile resistance

Phenotypically, leaf strength was significantly and strongly associated with all three morphological traits. Leaf strength positively covaried with LMA (r_P_ = 0.682 ± 0.034, p < 0.001) and leaf thickness (r_P_ = 0.738 ± 0.032, p < 0.001) but negatively covaried with leaf density (r_P_ = - 0.200 ± 0.046, p < 0.001). Leaf toughness was also significantly associated with these three morphological traits, but the associations were not as strong. As with leaf strength, leaf toughness positively covaried with LMA (r_P_ = 0.489 ± 0.041, p < 0.001) and leaf thickness (r_P_ = 0.523 ± 0.040, p < 0.001) and negatively covaried with leaf density (r_P_ = −0.124 ± 0.047, p < 0.01).

The genetic correlations between these traits show a similar pattern to the phenotypic correlations (Fig. 4). There were positive genetic correlations between LMA and leaf strength (r_G_ = 0.453 ± 0.076, p < 0.001) and LMA and leaf toughness (r_G_ = 0.403 ± 0.109, p < 0.001). In addition, leaf thickness was positively genetically correlated with leaf strength (r_G_ = 0.599 ± 0.071, p < 0.001) and leaf toughness (r_G_ = 0.507 ± 0.098, p < 0.001). There were marginally significant negative genetic correlations between leaf strength and leaf density (r_G_ = −0.188 ± 0.103, p = 0.055) and leaf toughness and leaf density (r_G_ = −0.216 ± 0.134, p = 0.10). Together, these results indicate that leaf morphology and composition contribute to leaf tensile resistance.

**Fig. 4.**
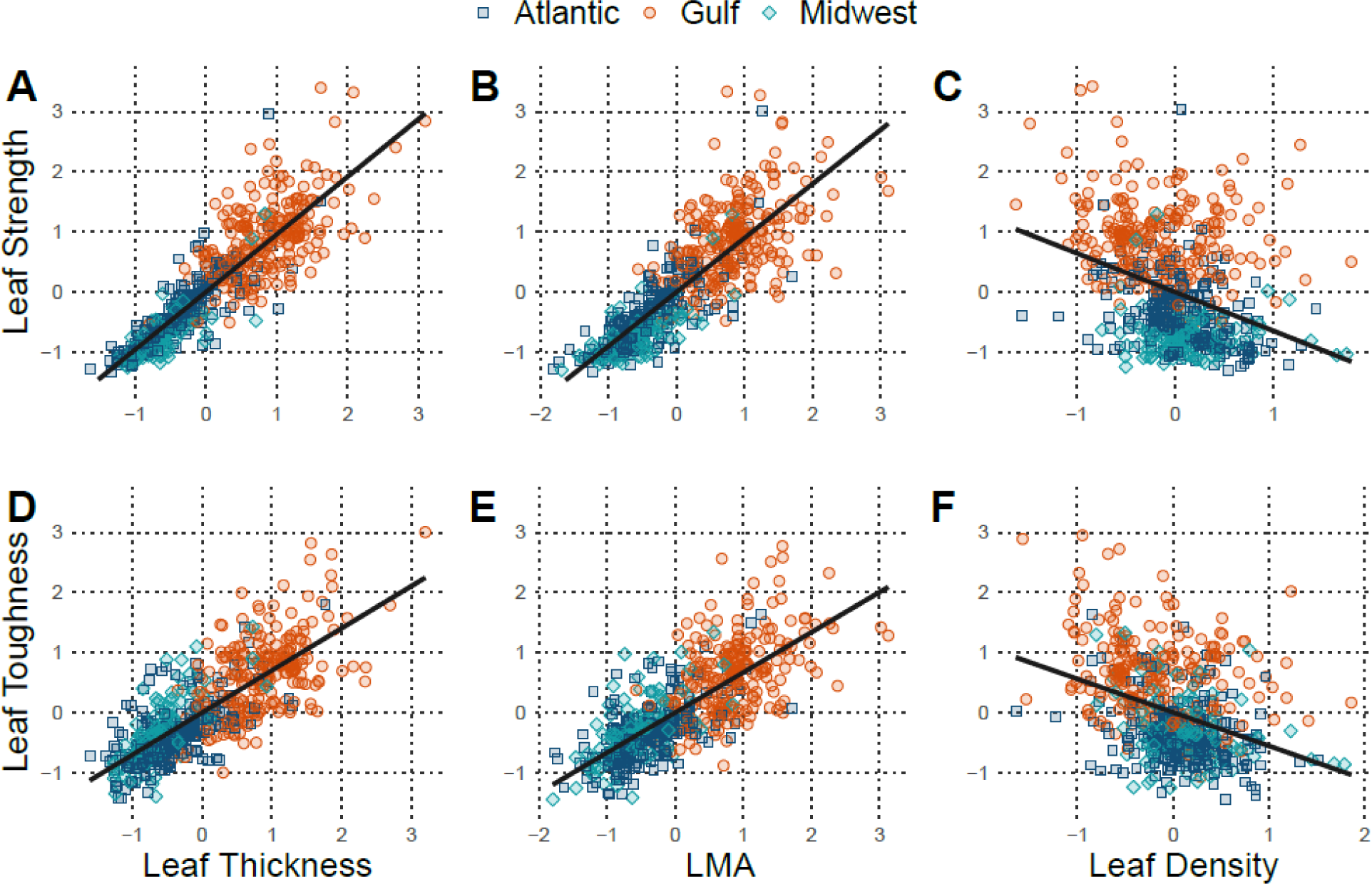
Genetic correlations between leaf structural resistance traits (strength and toughness) and LMA, leaf thickness, and leaf density. Models were fit using all genotypes; colors and shapes are added to differentiate subpopulations. Trend lines represent the fitted linear relationship between genetic best linear unbiased predictions (GBLUPs).

### Genome-wide association and candidate genes underlying leaf traits

Using GWA, we identified several variants associated with leaf tensile resistance and leaf morphology traits. Five significant single nucleotide polymorphisms (SNPs) were associated with leaf strength when using a Bonferroni-adjusted significance threshold (α = 0.05). At the more relaxed 5% false discovery rate significance threshold, 33 SNPs were associated with leaf strength across 11 chromosomes (Table S5; Fig. 5). There were 30 genes within 10 kilobase pairs of these SNPs (Table S6). Leaf toughness was not significantly associated with any variants at either the 5% false discovery rate threshold or the Bonferroni-adjusted threshold (Fig. S3A). Leaf morphological traits exhibited large differences in their degree of genetic association. Leaf area was associated with 292 variants across all 18 chromosomes at a 5% false discovery rate significance threshold (Table S5, Fig. S3B); 293 genes were located within 10 kilobase pairs of these SNPs (Table S6). Leaf lamina thickness, on the other hand, was only associated with one variant (Table S5, Fig. S3C), which was located near two genes (Table S6). LMA was not significantly associated with any variants (Fig. S3D).

**Fig. 5.**
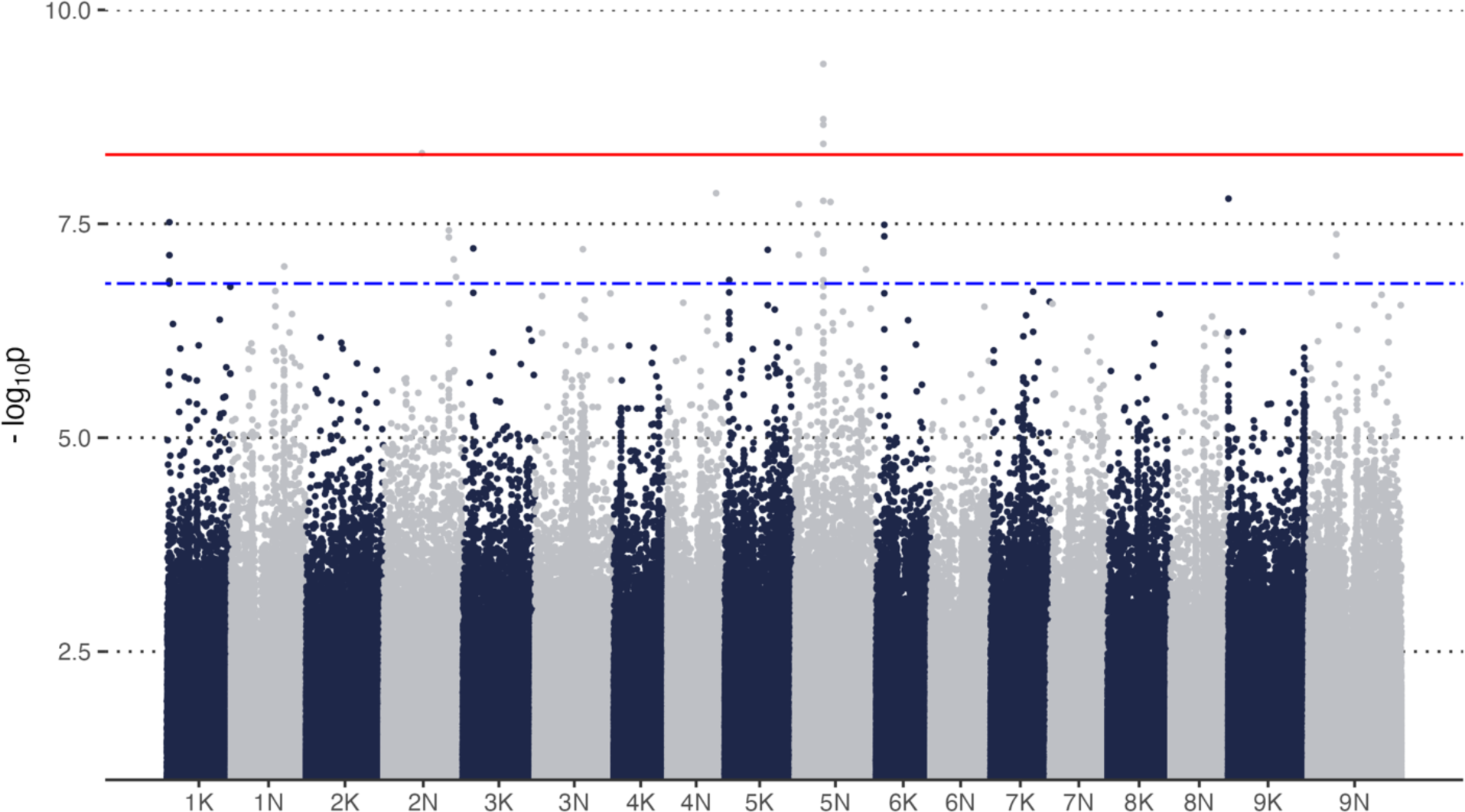
Manhattan plot showing the association between single nucleotide polymorphisms (SNPs) and leaf strength. The solid red line represents a genome-wide Bonferroni-adjusted significance threshold and the dashed blue line below represents a genome-wide 5% false discovery rate threshold; SNPs above these lines have a significant statistical association with leaf strength (−log_10_ *P*-value). For plotting purposes, we removed all SNPs whose −log_10_ p-value < 1.

When we looked for overlap in SNPs associated with leaf tensile resistance and leaf morphology using Cauchy combination tests, SNPs fell into four categories: SNPs associated with leaf area and leaf tensile resistance (n = 2), SNPs associated with leaf tensile resistance and multiple leaf morphological traits (n = 3), SNPs only associated with leaf strength (n = 30), and SNPs only associated with leaf area (n = 290). There was also a single SNP that was associated with leaf tensile resistance and leaf thickness; this was the same significant SNP that was associated with leaf thickness.

Finally, there were many candidate genes present within 10 kilobase pairs of significant SNPs in both univariate and combination test GWA. Specifically, five candidate genes were exclusively associated with leaf strength. These included genes related to stress responses, such as an ortholog of the regulatory gene *GF14* (Pavir.1NG310300; Chen et al. 2006), abscisic acid (Pavir.3KG137400; APA1; Sebastián et al. 2020), and gibberellic acid (Pavir.1NG310319; SLY1; Ariizumi et al. 2011). There were also genes associated with growth and development, including an ortholog of an auxin regulator *ABCB4* (Pavir.3KG136954; Kubeš et al. 2012) and an ortholog of *Cyp40* (Pavir.4NG131800), which is necessary for vegetative phase change in *Arabidopsis thaliana* (Berardini et al. 2001). We detected one candidate pleiotropic gene (Pavir.7NG287700) associated with leaf area and leaf tensile resistance—an ortholog of *AGO1* which is involved in auxin regulation (Sorin et al. 2005) and vegetative phase change (Yang et al. 2006) as well as normal leaf morphology (Bohmert et al. 1998, Jover-Gil et al. 2012). We also found a candidate pleiotropic gene (Pavir. 9KG134601) associated with leaf thickness and leaf tensile resistance, *NaPRT1*, which is involved in stress response (Hashida et al. 2009) and leaf senescence, the final stage of leaf development (Schippers et al. 2008, Hashida et al. 2009, Wu et al. 2016). These candidates suggest that genes related to stress response, auxin regulation within the cell, and development are critical in determining leaf tensile resistance and likely have pleiotropic effects on the leaf as a whole.

### Leaf tensile resistance increases with growing season length

Growing season length may be an important determinant of leaf tensile resistance, possibly resulting from changes in leaf morphology. We evaluated this hypothesis using mediation analysis, an approach that assesses the importance of indirect effects on a response. Overall, the effect of habitat-of-origin growing degree days on leaf strength and leaf toughness were both mediated by LMA (standardized effect on leaf strength = 0.1n82, p < 0.001; standardized effect on leaf toughness = 0.149, p < 0.001) and by leaf thickness (standardized effect on leaf strength = 0.241, p < 0.001; standardized effect on leaf toughness = 0.198, p < 0.001; Table S7, S8; Fig. 6). Moreover, there was a significant direct effect of growing degree days on leaf strength (standardized effect = 0.357, p < 0.001) and on leaf toughness (standardized effect = 0.174, p = 0.007). These large direct effects indicate that another important mediator is probably absent from our analysis.

**Fig. 6.**
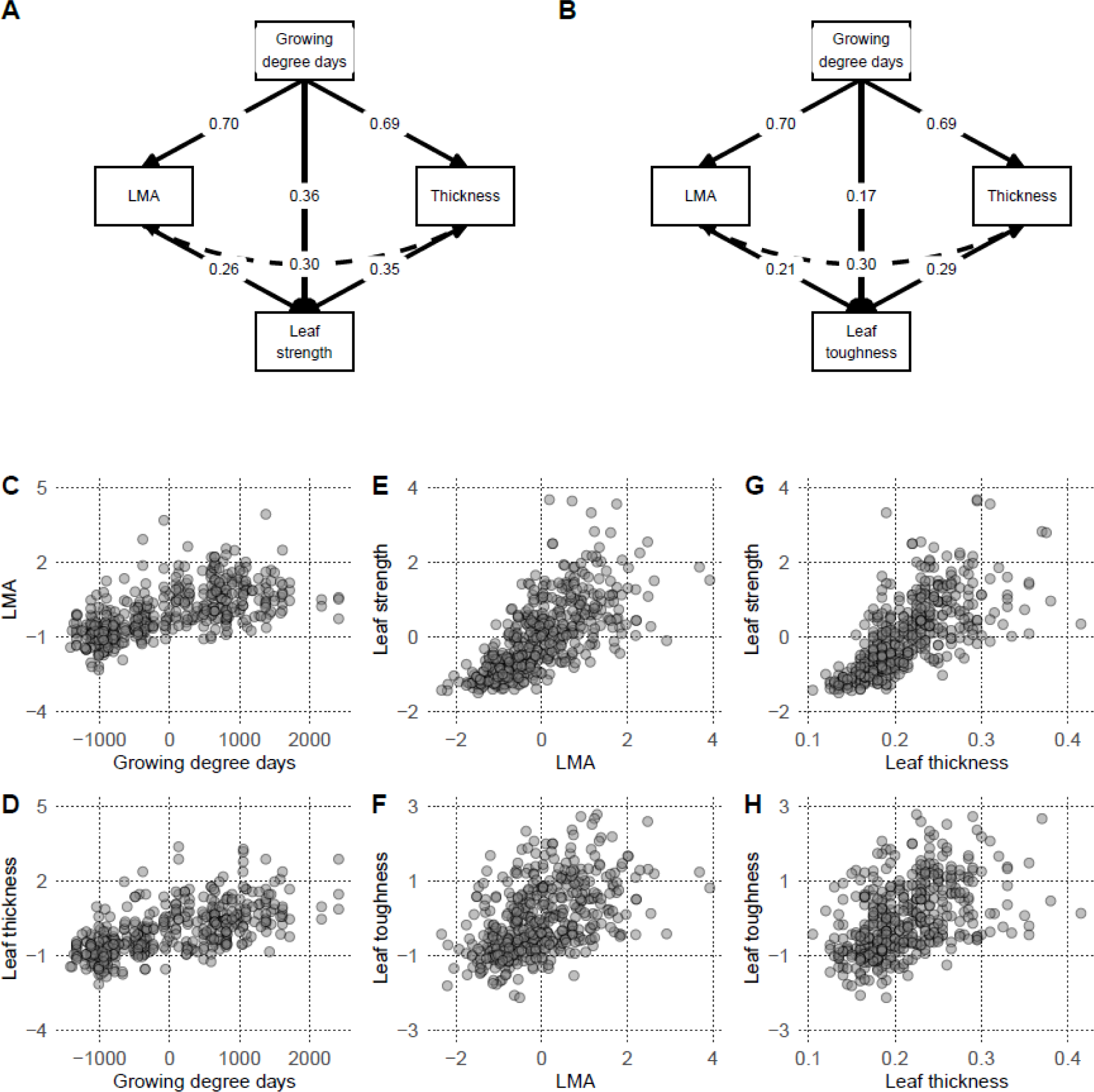
Direct and indirect effects of growing degree days, a proxy for growing season length, on A) leaf strength and B) leaf toughness showing the importance of two leaf morphological traits, LMA and leaf thickness. All coefficients are standardized; black lines denote that all model paths are significant. Panels C–G show bivariate relationships represented in panels A) and B). Growing degree days was mean-centered. All other variables were standardized such that across all plants, mean = 0 and standard deviation = 1.

Another possible explanation is that the effects of habitat-of-origin growing degree days on leaf strength and leaf toughness were mediated to differing degrees by LMA and leaf thickness in each subpopulation (Table S9, S10, Fig. S4). In the Atlantic and Gulf subpopulations the leaf strength effect was only significantly mediated by leaf thickness (Atlantic standardized effect = 0.149, p = 0.002; Gulf standardized effect = 0.084, p = 0.056), while the leaf strength effect in the Midwest subpopulation was mediated by both LMA and leaf thickness (LMA standardized effect = 0.114, p = 0.013; Thickness standardized effect = 0.108, p = 0.057; Table S9 Fig. S4A, S5). Moreover, the direct effect of growing degree days on leaf strength was only significant in the Atlantic subpopulation (standardized effect = 0.396, p < 0.001; Table S9, Fig. S4A).

The indirect effect of growing degree days on leaf toughness mediated by LMA and leaf thickness were also weak. In the Atlantic subpopulation, neither indirect effect was significant; in the Gulf and Midwest subpopulations, the indirect leaf thickness effect was marginally significant (Gulf standardized effect = 0.073, p = 0.081; Midwest standardized effect = 0.142, p = 0.087), but the indirect LMA effect was not significant (Table S10, Fig. S4B). Only in the Atlantic subpopulation was the direct effect of growing degree days on leaf toughness marginally significant (standardized effect = 0.193, p = 0.080; Table S10, Fig. S4B); in the other two subpopulations, that direct effect was highly non-significant (p > 0.75). Together, these results demonstrate that leaf morphology (leaf thickness and LMA) contributes substantially to the effect of growing season length on leaf tensile resistance (i.e., leaf morphology mediates this effect) across all genotypes of *P. virgatum*. Importantly, though, within genetic subpopulations, these relationships are more consistently mediated by leaf thickness than LMA.

## Discussion

In this study, we explored the intraspecific phenotypic and genetic variation in leaf tensile resistance in *P. virgatum*. We found that the three genetic subpopulations of *P. virgatum* exhibited large phenotypic differences in leaf tensile resistance, that leaf tensile resistance was highly heritable within and across subpopulations, and that high leaf tensile resistance was associated with high fitness as measured by biomass production. Moreover, leaf morphological traits, especially LMA and leaf thickness were positively correlated—both phenotypically and genetically—with leaf tensile resistance and these traits partially mediated the positive effect of growing season length on leaf tensile resistance. Additionally, multiple candidate genes for leaf tensile resistance were related to stress responses, auxin regulation within the cell, and leaf development and morphology; some of these candidate genes may have pleiotropic effects. Together, these results suggest that differences in climate across the broad range of *P. virgatum* contribute to the differences in leaf tensile resistance observed in this study—plants from the southernmost *P. virgatum* subpopulation, where growing seasons are longer, have evolved to produce stronger leaves with morphological traits (i.e., high LMA and leaf thickness) that may enhance leaf longevity. These results further suggest that leaf tensile resistance in *P. virgatum* may evolve in response to current and future climate change.

Much of the natural variation in leaf tensile resistance in *P. virgatum* can be attributed to the genetic variation present within the species. The high heritability (h^2^) of leaf strength and leaf toughness in *P. virgatum* indicate that genetics is critical in determining leaf structural resistance. This is consistent with a study on stem structural resistance in *Sorghum bicolor*, which found a significant genetic contribution to stem strength, stiffness, and rigidity (Gomez et al. 2020). Additionally, we identified several candidate genes associated with leaf strength, including genes functioning in leaf development, responses to abiotic stress, and hormones like abscisic acid, auxin, and gibberellic acid. Unlike a similar study in *A. thaliana*, which demonstrated an important role for cell wall composition in leaf strength (Balsamo et al. 2015), we did not detect any candidate genes associated cell wall composition. Our lack of evidence here could have resulted from low statistical signal—characteristics like leaf tensile resistance may be highly polygenic and our sample size was relatively small, which would limit our ability to detect genes of moderate effect. In support of this statistical argument, we identified a candidate gene associated with cell wall composition at the less stringent 10% false discovery rate. It is likely that across tissues and systems, mechanical properties have a genetic basis which implies that structurally robust plants can be bred in agricultural systems and tissue toughness in many naturally occurring plant populations may adapt to environmental conditions. Specifically, in agricultural settings, stem lodging (i.e., stem bending near ground level) can reduce plant productivity and crop yields (Foulkes et al. 2011); in biofuel crops, stronger leaves are more likely to remain attached to the stem and contribute to biomass yield (Karp and Shield 2008). In natural systems, this means that in many species tissue toughness will be related to biotic pressures, such as herbivory, or abiotic pressures (Pérez-Harguindeguy et al. 2003, Moles et al. 2011).

Although there are numerous studies on the drivers of leaf toughness across species (Kitajima and Poorter 2010, Onoda et al. 2017), less research has been done within individual species. Because interspecific and intraspecific patterns do not always concur (Agrawal 2020), it is worthwhile to investigate the intraspecific relationship between leaf morphology and leaf structural resistance. We found that leaf strength and toughness were both positively phenotypically correlated with LMA and leaf thickness. This result is consistent with the findings of interspecific studies (Onoda et al. 2011). On the other hand, leaf strength and toughness were marginally negatively correlated with leaf density. According to Onoda et al. (2011), we would expect a positive relationship between leaf structural resistance and leaf density. Similarly, a study amongst oaks (*Quercus spp.*) found that LMA and leaf thickness were highly correlated with punch strength, a measure of resistance to puncturing, while there was a weaker relationship between leaf density and punch strength (Alonso-Forn et al. 2023). Although density may be a useful predictor of leaf structural resistance across a large phylogenetic scale, LMA and leaf thickness may be more reliable predictors and likely have a more direct contribution to leaf structural resistance at the intraspecific level. In addition, intraspecific studies give us the opportunity to explore genetic relationships between traits. We found a significant genetic relationships between leaf tensile resistance and both LMA and leaf thickness, including two candidate pleiotropic genes (orthologs of *AGO1* and *NaPRT1;* Pavir.7NG287700 and Pavir. 9KG134601, respectively) that may affect both leaf tensile resistance and morphology. Together, the results of these genetic analyses indicate that there is an underlying phenotypic and genetic relationship between leaf morphology and leaf tensile resistance.

Even though our findings show that genetics and leaf morphology strongly influence leaf tensile resistance, there were some limitations. First, population structure can be a confounding factor for quantitative genetic and genomic analyses (Sul et al. 2018). The high heritability of leaf strength and leaf toughness within the Atlantic and Gulf subpopulations, however, suggests that strong population structure was not the primary driver of these results. Second, only one study site was used, which prevents us from knowing how plasticity or genotype-by-environment interactions affect leaf tensile resistance in *P. virgatum* (Wilson et al. 2010). The conditions at this site differed from the climate-of-origin of some of plants, particularly those from the Midwest and Atlantic subpopulations, which may result in changes to their observed phenotypes (Johnson et al. 2022, Napier et al. 2023). Third, we sampled considerably fewer Midwest plants (90) than Gulf and Atlantic plants (201 and 164, respectively). The combined effect of fewer Midwest plants being grown and measured in conditions vastly different from their natural habitat could have contributed to the low heritability observed in the Midwest subpopulation. Despite these limitations, the sample of genotypes still had sufficient diversity and is likely representative of the natural genetic and phenotypic variation in *P. virgatum*.

In addition to having sufficient variation in leaf tensile resistance to adapt to different environments, there is evidence that *P. virgatum* has evolved tougher leaves in response to longer growing seasons across and within subpopulations. The Midwest and the Atlantic subpopulations’ ecological strategies may favor a faster growth rate to make the most of a shorter growing season, while the longer Gulf subpopulation growing seasons may promote the development of stronger, but more resource-expensive leaves (Wright et al. 2004, Onoda et al. 2011, Lowry et al. 2014, Heckman et al. 2020, Wang et al. 2023). At this site, which is in the southern portion of the *P. virgatum* range, and thus has a relatively long growing season, the individuals with tougher leaves had higher aboveground biomass, an indicator of higher fitness (Lowry et al. 2019), than those with weaker leaves. The high variation in leaf structural resistance present within *P. virgatum* and its strong links with the climate of genotype origin suggest that structural resistance may be an important contributor to the success of genotypes under climate change, potentially promoting local adaptation to changing growing season length.

## Conclusions

Despite the potential costs associated with high leaf structural resistance, plants in this study with tougher and stronger leaves produced more biomass—a proxy for fitness—which may be associated with the long growing season at our common garden. Leaf tensile resistance was strongly associated with leaf morphological traits like leaf thickness and LMA, although these relationships often differed among genetic subpopulations. This suggests that leaves may be responding to different selective pressures across the range of *P. virgatum*. Together, the high heritability of leaf tensile resistance, the strong genetic links between tensile resistance and leaf morphology, and the impact of growing season length on tensile resistance suggest that leaves of *P. virgatum* will be able to respond to a warming climate in the coming decades.

## Acknowledgements

Thank you to Angelo Filonzi for making the tensile resistance assays possible. Thank you to Jason Bonnette and Emily Breckenridge for helping with data collection. Thank you to David Lowry, Acer VanWallendael, Daniel Anstett, Joseph Edwards, Otto Kailing, Leslie Kollar, Jason Olsen, Madison Plunkert, and Lauren Stanley for providing comments and suggestions. The findings and conclusions of this publication are those of the authors and should not be construed to represent any official USDA or U.S. Government determination or policy.

## Author contributions

TEJ, AB, and RWH conceived the study; PCD and RWH implemented the experiment and analyzed the data; PCD and RWH wrote the manuscript with contributions from TEJ and AB.

## References

Agrawal, A. A. 2020. A scale-dependent framework for trade-offs, syndromes, and specialization in organismal biology. Ecology 101:e02924.

Agrawal, A. A., and M. Fishbein. 2006. Plant defense syndromes. Ecology 87:S132–S149.

Alonso-Forn, D., D. Sancho-Knapik, M. D. Fariñas, M. Nadal, R. Martín-Sánchez, J. P. Ferrio, … E. Gil-Pelegrín. 2023. Disentangling leaf structural and material properties in relationship to their anatomical and chemical compositional traits in oaks (*Quercus* L.). Annals of Botany 131:789–800.

Anstett, D. N., K. A. Nunes, C. Baskett, and P. M. Kotanen. 2016. Sources of controversy surrounding latitudinal patterns in herbivory and defense. Trends in Ecology & Evolution 31:789–802.

Ariizumi, T., P. K. Lawrence, and C. M. Steber. 2011. The role of two F-box proteins, *SLEEPY1* and *SNEEZY*, in *Arabidopsis* gibberellin signaling. Plant Physiology 155:765–775.

Balsamo, R., M. Boak, K. Nagle, B. Peethambaran, and B. Layton. 2015. Leaf biomechanical properties in *Arabidopsis thaliana* polysaccharide mutants affect drought survival. Journal of Biomechanics 48:4124–4129.

Berardini, T. Z., K. Bollman, H. Sun, and R. Scott Poethig. 2001. Regulation of vegetative phase change in *Arabidopsis thaliana* by cyclophilin 40. Science 291:2405–2407.

Bohmert, K., I. Camus, C. Bellini, D. Bouchez, M. Caboche, and C. Benning. 1998. *AGO1* defines a novel locus of *Arabidopsis* controlling leaf development. The EMBO Journal 17:170–180.

Butler, D. 2021. asreml: fits the linear mixed model. VSN International.

Casler, M. D. 2012. Switchgrass Breeding, Genetics, and Genomics. Pages 29-53 *in* A. Monti, editor. Switchgrass: A Valuable Biomass Crop for Energy. Springer London, London.

Casler, M. D., C. M. Tobias, S. M. Kaeppler, C. R. Buell, Z.-Y. Wang, P. Cao, … P. Ronald. 2011. The switchgrass genome: tools and strategies. The Plant Genome 4.

Chen, F., Q. Li, L. Sun, and Z. He. 2006. The rice 14-3-3 gene family and its involvement in responses to biotic and abiotic stress. DNA Research 13:53–63.

Coley, P. D., and J. A. Barone. 1996. Herbivory and plant defenses in tropical forests. Annual review of Ecology and Systematics 27:305–335.

Coley, P. D., J. P. Bryant, and F. S. Chapin. 1985. Resource availability and plant antiherbivore defense. Science 230:895–899.

Foulkes, M. J., G. A. Slafer, W. J. Davies, P. M. Berry, R. Sylvester-Bradley, P. Martre, … M. P. Reynolds. 2011. Raising yield potential of wheat. III. Optimizing partitioning to grain while maintaining lodging resistance. Journal of Experimental Botany 62:469–486.

Franklin, O. D., and M. B. Morrissey. 2017. Inference of selection gradients using performance measures as fitness proxies. Methods in Ecology and Evolution 8:663–677.

Gomez, F. E., J. E. Mullet, A. H. Muliana, K. J. Niklas, and W. L. Rooney. 2020. The genetic architecture of biomechanical traits in sorghum. Crop Science 60:82–99.

Grady, K. C., D. C. Laughlin, S. M. Ferrier, T. E. Kolb, S. C. Hart, G. J. Allan, and T. G. Whitham. 2013. Conservative leaf economic traits correlate with fast growth of genotypes of a foundation riparian species near the thermal maximum extent of its geographic range. Functional Ecology 27:428–438.

Hales, T. C., and C. F. Miniat. 2017. Soil moisture causes dynamic adjustments to root reinforcement that reduce slope stability. Earth Surface Processes and Landforms 42:803–813.

Hashida, S.-n., H. Takahashi, and H. Uchimiya. 2009. The role of NAD biosynthesis in plant development and stress responses. Annals of Botany 103:819–824.

Heckman, R. W., A. R. Khasanova, N. S. Johnson, S. Weber, J. E. Bonnette, M. J. Aspinwall, … C. V. Hawkes. 2020. Plant biomass, not plant economics traits, determines responses of soil CO_2_ efflux to precipitation in the C_4_ grass *Panicum virgatum*. Journal of Ecology 108:2095–2106.

Johnson, L. C., M. B. Galliart, J. D. Alsdurf, B. R. Maricle, S. G. Baer, N. M. Bello, … A. B. Smith. 2022. Reciprocal transplant gardens as gold standard to detect local adaptation in grassland species: New opportunities moving into the 21st century. Journal of Ecology 110:1054–1071.

Jover-Gil, S., H. Candela, P. Robles, V. Aguilera, J. M. Barrero, J. L. Micol, and M. R. Ponce. 2012. The microRNA pathway genes *AGO1*, *HEN1* and *HYL1* participate in leaf proximal–distal, venation and stomatal patterning in *Arabidopsis*. Plant and Cell Physiology 53:1322–1333.

Karp, A., and I. Shield. 2008. Bioenergy from plants and the sustainable yield challenge. New Phytologist 179:15–32.

Kikuzawa, K., Y. Onoda, I. J. Wright, and P. B. Reich. 2013. Mechanisms underlying global temperature-related patterns in leaf longevity. Global Ecology and Biogeography 22:982–993.

Kitajima, K., and L. Poorter. 2010. Tissue-level leaf toughness, but not lamina thickness, predicts sapling leaf lifespan and shade tolerance of tropical tree species. New Phytologist 186:708–721.

Kremling, K. A. G., C. H. Diepenbrock, M. A. Gore, E. S. Buckler, and N. B. Bandillo. 2019. Transcriptome-wide association supplements genome-wide association in *Zea mays*. G3 Genes|Genomes|Genetics 9:3023–3033.

Kubeš, M., H. Yang, G. L. Richter, Y. Cheng, E. Młodzińska, X. Wang, … A. S. Murphy. 2012. The *Arabidopsis* concentration-dependent influx/efflux transporter *ABCB4* regulates cellular auxin levels in the root epidermis. The Plant Journal 69:640–654.

Kudo, G. 1996. Intraspecific variation of leaf traits in several deciduous species in relation to length of growing season. Écoscience 3:483–489.

Lande, R., and S. J. Arnold. 1983. The measurement of selection on correlated characters. Evolution:1210–1226.

Lenth, R. 2023. emmeans: Estimated Marginal Means, aka Least-Squares Means. https://CRAN.R-project.org/package=emmeans.

Li, H., and R. Durbin. 2009. Fast and accurate short read alignment with Burrows–Wheeler transform. Bioinformatics 25:1754–1760.

Liu, Y. 2020. ACAT: Aggregaged Cauchy Associaion Test (ACAT).

Lovell, J. T., A. H. MacQueen, S. Mamidi, J. Bonnette, J. Jenkins, J. D. Napier, … J. Schmutz. 2021. Genomic mechanisms of climate adaptation in polyploid bioenergy switchgrass. Nature 590:438–444.

Lowry, D. B., K. D. Behrman, P. Grabowski, G. P. Morris, J. R. Kiniry, and T. E. Juenger. 2014. Adaptations between ecotypes and along environmental gradients in Panicum virgatum. American Naturalist 183:682–692.

Lowry, D. B., J. T. Lovell, L. Zhang, J. Bonnette, P. A. Fay, R. B. Mitchell, … T. E. Juenger. 2019. QTL × environment interactions underlie adaptive divergence in switchgrass across a large latitudinal gradient. Proceedings of the National Academy of Sciences 116:12933.

Lucas, P. W., I. M. Turner, N. J. Dominy, and N. Yamashita. 2000. Mechanical defences to herbivory. Annals of Botany 86:913–920.

Lynch, M., and B. Walsh. 1998. Genetics and analysis of quantitative traits. Sinauer, Sunderland, MA.

Mackay, T. F. C. 2001. The genetic architecture of quantitative traits. Annual Review of Genetics 35:303–339.

Mason, C. M., and L. A. Donovan. 2015. Evolution of the leaf economics spectrum in herbs: Evidence from environmental divergences in leaf physiology across *Helianthus* (Asteraceae). Evolution 69:2705–2720.

Moles, A. T., I. R. Wallis, W. J. Foley, D. I. Warton, J. C. Stegen, A. J. Bisigato, … L. D. Prior. 2011. Putting plant resistance traits on the map: a test of the idea that plants are better defended at lower latitudes. New Phytologist 191:777–788.

Napier, J. D., R. W. Heckman, and T. E. Juenger. 2023. Gene-by-environment interactions in plants: Molecular mechanisms, environmental drivers, and adaptive plasticity. The Plant Cell 35:109–124.

Nardini, A. 2022. Hard and tough: the coordination between leaf mechanical resistance and drought tolerance. Flora 288:152023.

Onoda, Y., M. Westoby, P. B. Adler, A. M. F. Choong, F. J. Clissold, J. H. C. Cornelissen, … N. Yamashita. 2011. Global patterns of leaf mechanical properties. Ecology Letters 14:301–312.

Onoda, Y., I. J. Wright, J. R. Evans, K. Hikosaka, K. Kitajima, Ü. Niinemets, … M. Westoby. 2017. Physiological and structural tradeoffs underlying the leaf economics spectrum. New Phytologist 214:1447–1463.

Palik, D. J., A. A. Snow, A. L. Stottlemyer, M. N. Miriti, and E. A. Heaton. 2016. Relative performance of non-local cultivars and local, wild populations of Switchgrass (*Panicum virgatum*) in competition experiments. PLOS ONE 11:e0154444.

Pérez-Harguindeguy, N., S. Díaz, F. Vendramini, J. H. C. Cornelissen, D. E. Gurvich, and M. Cabido. 2003. Leaf traits and herbivore selection in the field and in cafeteria experiments. Austral Ecology 28:642–650.

Pinheiro, J., D. Bates, and R. C. Team. 2023. nlme: Linear and Nonlinear Mixed Effects Models. https://CRAN.R-project.org/package=nlme.

Pinheiro, J., and D. M. Bates. 2000. Mixed-Effects Models in S and S-PLUS. Springer, New York.

Privé, F., H. Aschard, A. Ziyatdinov, and M. G. B. Blum. 2018. Efficient analysis of large-scale genome-wide data with two R packages: bigstatsr and bigsnpr. Bioinformatics 34:2781–2787.

Rosseel, Y. 2012. lavaan: An R package for structural equation modeling. Journal of Statistical Software 48:1–36.

Schippers, J. H. M., A. Nunes-Nesi, R. Apetrei, J. Hille, A. R. Fernie, and P. P. Dijkwel. 2008. The *Arabidopsis onset of leaf death5* mutation of quinolinate synthase affects nicotinamide adenine dinucleotide biosynthesis and causes early ageing. The Plant Cell 20:2909–2925.

Sebastián, D. I., F. D. Fernando, D. G. Raúl, and G. M. Gabriela. 2020. Overexpression of *Arabidopsis* aspartic protease *APA1* gene confers drought tolerance. Plant Science 292:110406.

Sorin, C. l., J. D. Bussell, I. Camus, K. Ljung, M. Kowalczyk, G. Geiss, … C. Bellini. 2005. Auxin and light control of adventitious rooting in *Arabidopsis* require *ARGONAUTE1*. The Plant Cell 17:1343–1359.

Sparks, A. H. 2018. nasapower: a NASA POWER global meteorology, surface solar energy and climatology data client for R. Journal of Open Source Software 3:1035.

Sparks, A. H. 2023. nasapower: NASA-POWER Data from R. https://CRAN.R-project.org/package=nasapower.

Stinchcombe, J. R., A. F. Agrawal, P. A. Hohenlohe, S. J. Arnold, and M. W. Blows. 2008. Estimating nonlinear selection gradients using quadratic regression coefficients: double or nothing? Evolution 62:2435–2440.

Storey, J. D., and R. Tibshirani. 2003. Statistical significance for genomewide studies. Proceedings of the National Academy of Sciences 100:9440–9445.

Sul, J. H., L. S. Martin, and E. Eskin. 2018. Population structure in genetic studies: Confounding factors and mixed models. PLOS Genetics 14:e1007309.

VanRaden, P. M. 2008. Efficient methods to compute genomic predictions. Journal of Dairy Science 91:4414–4423.

Walsh, B., and M. W. Blows. 2009. Abundant genetic variation + strong selection = multivariate genetic constraints: a geometric view of adaptation. Annual Review of Ecology, Evolution, and Systematics 40:41–59.

Wang, H., I. C. Prentice, I. J. Wright, D. I. Warton, S. Qiao, X. Xu, … N. C. Stenseth. 2023. Leaf economics fundamentals explained by optimality principles. Science Advances 9:eadd5667.

Westbrook, J. W., K. Kitajima, J. G. Burleigh, W. J. Kress, D. L. Erickson, and S. J. Wright. 2011. What makes a leaf tough? patterns of correlated evolution between leaf toughness traits and demographic rates among 197 shade-tolerant woody species in a neotropical forest. The American Naturalist 177:800–811.

Wilson, A. J., D. Réale, M. N. Clements, M. M. Morrissey, E. Postma, C. A. Walling, … D. H. Nussey. 2010. An ecologist’s guide to the animal model. Journal of Animal Ecology 79:13–26.

Wright, I. J., P. B. Reich, M. Westoby, D. D. Ackerly, Z. Baruch, F. Bongers, … R. Villar. 2004. The worldwide leaf economics spectrum. Nature 428:821–827.

Wu, D., X. Li, R. Tanaka, J. C. Wood, L. E. Tibbs-Cortes, M. Magallanes-Lundback, … M. A. Gore. 2022. Combining GWAS and TWAS to identify candidate causal genes for tocochromanol levels in maize grain. Genetics 221.

Wu, L., D. Ren, S. Hu, G. Li, G. Dong, L. Jiang, … L. Guo. 2016. Mutation of *OsNaPRT1* in the NAD salvage pathway leads to withered leaf tips in rice. Plant Physiology 171:pp.01898.02015-pp.01898.02015.

Yang, L., W. Huang, H. Wang, R. Cai, Y. Xu, and H. Huang. 2006. Characterizations of a hypomorphic *Argonaute1* mutant reveal novel *AGO1* functions in *Arabidopsis* lateral organ development. Plant Molecular Biology 61:63–78.

Yu, J., G. Pressoir, W. H. Briggs, I. Vroh Bi, M. Yamasaki, J. F. Doebley, … E. S. Buckler. 2006. A unified mixed-model method for association mapping that accounts for multiple levels of relatedness. Nature Genetics 38:203–208.

Zhao, X., J. J. G. Lynch, and Q. Chen. 2010. Reconsidering Baron and Kenny: myths and truths about mediation analysis. Journal of Consumer Research 37:197–206.

Züst, T., and A. A. Agrawal. 2017. Trade-offs between plant growth and defense against insect herbivory: an emerging mechanistic synthesis. Annual Review of Plant Biology 68:513–534.

