## Supplemental Tables and Figures for "The shared genetic basis of leaf morphology and tensile resistance underlies the effect of growing season length in a widespread perennial grass"

**Table S1** ANOVA table showing effects of A) leaf strength and B) leaf toughness on plant relative biomass

| | Parameter | DF | F | *P* |
|---|---|---|---|---|
| A) Leaf strength | Intercept | 1, 434 | 1347.69 | < 0.001 |
| | Leaf strength | 1, 434 | 74.93 | < 0.001 |
| | Leaf strength$^2$ | 1, 434 | 20.93 | < 0.001 |
| | Subpopulation | 2, 434 | 0 | 0.999 |
| | Leaf strength × Subpopulation | 2, 434 | 24.44 | < 0.001 |
| | Leaf strength$^2$ × Subpopulation | 2, 434 | 1.95 | 0.144 |
| B) Leaf toughness | Intercept | 1, 434 | 1084.24 | < 0.001 |
| | Leaf toughness | 1, 434 | 13.73 | < 0.001 |
| | Leaf toughness$^2$ | 1, 434 | 12.41 | < 0.001 |
| | Subpopulation | 2, 434 | 0 | 0.999 |
| | Leaf toughness × Subpopulation | 2, 434 | 3.56 | 0.029 |
| | Leaf toughness$^2$ × Subpopulation | 2, 434 | 0.73 | 0.482 |

**Table S2** Table showing standardized effects of A) leaf strength and B) leaf toughness on plant relative biomass. Estimate is parameter estimate from model; all quadratic term estimates are doubled.

| | Parameter | Value | Std Error |
|---|---|---|---|
| A) Leaf strength | Intercept | 1.104 | 0.049 |
| | Leaf strength | 0.594 | 0.055 |
| | Leaf strength$^2$ | -0.221 | 0.042 |
| | SubpopulationGulf | -0.025 | 0.070 |
| | SubpopulationMidwest | -0.063 | 0.083 |
| | Leaf strength × SubpopulationGulf | -0.473 | 0.072 |
| | Leaf strength × SubpopulationMidwest | -0.254 | 0.100 |
| | Leaf strength$^2$ × SubpopulationGulf | 0.060 | 0.068 |
| | Leaf strength$^2$ × SubpopulationMidwest | 0.139 | 0.071 |
| B) Leaf toughness | Intercept | 1.123 | 0.064 |
| | Leaf toughness | 0.288 | 0.060 |
| | Leaf toughness$^2$ | -0.239 | 0.078 |
| | SubpopulationGulf | -0.066 | 0.086 |
| | SubpopulationMidwest | -0.034 | 0.105 |
| | Leaf toughness × SubpopulationGulf | -0.224 | 0.078 |
| | Leaf toughness × SubpopulationMidwest | -0.082 | 0.096 |
| | Leaf toughness$^2$ × SubpopulationGulf | 0.126 | 0.104 |
| | Leaf toughness$^2$ × SubpopulationMidwest | 0.061 | 0.127 |

**Table S3** Trends in responses of leaf strength and leaf toughness to aboveground biomass, separately by population. *P* represents whether the trend differs significantly from 0.

| Response | Gradient | Subpopulation | Trend | Std Error | *P* |
|---|---|---|---|---|---|
| A) Leaf strength | Linear | Atlantic | 0.59 | 0.06 | < 0.001 |
| | | Gulf | 0.12 | 0.05 | 0.008 |
| | | Midwest | 0.34 | 0.08 | < 0.001 |
| | Quadratic | Atlantic | -0.22 | 0.04 | < 0.001 |
| | | Gulf | -0.16 | 0.05 | 0.003 |
| | | Midwest | -0.08 | 0.06 | 0.150 |
| B) Leaf toughness | Linear | Atlantic | 0.29 | 0.06 | < 0.001 |
| | | Gulf | 0.06 | 0.05 | 0.201 |
| | | Midwest | 0.21 | 0.08 | 0.006 |
| | Quadratic | Atlantic | -0.24 | 0.08 | 0.002 |
| | | Gulf | -0.11 | 0.07 | 0.102 |
| | | Midwest | -0.18 | 0.10 | 0.076 |

**Table S4** Post-hoc test results comparing the effects of leaf strength and leaf toughness on aboveground biomass between genetic subpopulations.

| Response | Gradient | Contrast | Estimate | Std Error | *P* |
|---|---|---|---|---|---|
| A) Leaf strength | Linear | Atlantic - Gulf | 0.47 | 0.07 | < 0.001 |
| | | Atlantic - Midwest | 0.25 | 0.10 | 0.032 |
| | | Gulf - Midwest | -0.22 | 0.10 | 0.056 |
| | Quadratic | Atlantic - Gulf | -0.06 | 0.07 | 0.649 |
| | | Atlantic - Midwest | -0.14 | 0.07 | 0.123 |
| | | Gulf - Midwest | -0.08 | 0.08 | 0.575 |
| B) Leaf toughness | Linear | Atlantic - Gulf | 0.22 | 0.08 | 0.012 |
| | | Atlantic - Midwest | 0.08 | 0.10 | 0.672 |
| | | Gulf - Midwest | -0.14 | 0.09 | 0.258 |
| | Quadratic | Atlantic - Gulf | -0.13 | 0.10 | 0.450 |
| | | Atlantic - Midwest | -0.06 | 0.13 | 0.883 |
| | | Gulf - Midwest | 0.07 | 0.12 | 0.854 |

**Table S7** Model parameters from mediation analysis of growing degree days on leaf strength. 95% CIs were calculated via 5000 bootstrapping iterations. The indirect LMA effect is the product of the LMA ~ GDD effect and the Leaf strength ~ LMA effect; the indirect thickness effect is the product of the Thickness ~ GDD effect and the Leaf strength ~ Thickness effect. The total effect is the sum of the two indirect effects and the direct Leaf strength ~ GDD effect. ~ indicates a hypothesized causal relationship; ~~ indicates a hypothesized non-causal relationship.

| **Path** | **Estimate** | **Std Error** | **95% CIs** | **z** | ***P*** |
|---|---|---|---|---|---|
| LMA ~ GDD | 0.699 | 0.043 | 0.616, 0.784 | 16.211 | < 0.001 |
| Thickness ~ GDD | 0.687 | 0.039 | 0.613, 0.764 | 17.644 | < 0.001 |
| Leaf strength ~ LMA | 0.261 | 0.049 | 0.166, 0.361 | 5.299 | < 0.001 |
| Leaf strength ~ Thickness | 0.351 | 0.059 | 0.238, 0.468 | 5.998 | < 0.001 |
| Leaf strength ~ GDD | 0.357 | 0.048 | 0.264, 0.454 | 7.366 | < 0.001 |
| LMA ~~ Thickness | 0.300 | 0.034 | 0.238, 0.374 | 8.881 | < 0.001 |
| LMA ~~ LMA | 0.623 | 0.059 | 0.520, 0.754 | 10.583 | < 0.001 |
| Thickness ~~ Thickness | 0.572 | 0.052 | 0.480, 0.690 | 10.917 | < 0.001 |
| Leaf strength ~~ Leaf strength | 0.375 | 0.042 | 0.303, 0.472 | 8.849 | < 0.001 |
| GDD ~~ GDD | 0.797 | 0 | 0.797, 0.797 | -- | -- |
| LMA ~1 | -0.028 | 0.038 | -0.101, 0.048 | -0.728 | 0.466 |
| Thickness ~1 | -0.040 | 0.037 | -0.110, 0.033 | -1.082 | 0.279 |
| Leaf strength ~1 | 0.002 | 0.030 | -0.056, 0.064 | 0.055 | 0.956 |
| GDD ~1 | 0.001 | 0 | 0.001, 0.001 | -- | -- |
| Indirect LMA effect | 0.182 | 0.036 | 0.115, 0.257 | 5.092 | < 0.001 |
| Indirect Thickness effect | 0.241 | 0.042 | 0.163, 0.328 | 5.726 | < 0.001 |
| Total effect | 0.781 | 0.049 | 0.264, 0.454 | 7.365 | < 0.001 |

**Table S9** Model parameters from mediation analysis of growing degree days on leaf toughness. 95% CIs were calculated via 5000 bootstrapping iterations. The indirect LMA effect is the product of the LMA ~ GDD effect and the Leaf toughness ~ LMA effect; the indirect thickness effect is the product of the Thickness ~ GDD effect and the Leaf toughness ~ Thickness effect. The total effect is the sum of the two indirect effects and the direct Leaf toughness ~ GDD effect. ~ indicates a hypothesized causal relationship; ~~ indicates a hypothesized non-causal relationship.

| **Path** | **Estimate** | **Std Error** | **95% CIs** | **z** | ***P*** |
|---|---|---|---|---|---|
| LMA ~ GDD | 0.699 | 0.043 | 0.615, 0.787 | 16.139 | < 0.001 |
| Thickness ~ GDD | 0.687 | 0.038 | 0.613, 0.764 | 17.871 | < 0.001 |
| Leaf toughness ~ LMA | 0.213 | 0.059 | 0.100, 0.329 | 3.629 | < 0.001 |
| Leaf toughness ~ Thickness | 0.289 | 0.072 | 0.151, 0.433 | 4.030 | < 0.001 |
| Leaf toughness ~ GDD | 0.174 | 0.064 | 0.048, 0.299 | 2.718 | 0.007 |
| LMA ~~ Thickness | 0.300 | 0.034 | 0.237, 0.370 | 8.940 | < 0.001 |
| LMA ~~ LMA | 0.623 | 0.058 | 0.526, 0.756 | 10.800 | < 0.001 |
| Thickness ~~ Thickness | 0.572 | 0.052 | 0.483, 0.688 | 11.040 | < 0.001 |
| Leaf toughness ~~ Leaf toughness | 0.686 | 0.049 | 0.602, 0.801 | 13.957 | < 0.001 |
| GDD ~~ GDD | 0.797 | 0 | 0.797, 0.797 | -- | -- |
| LMA ~1 | -0.028 | 0.038 | -0.102, 0.049 | -0.718 | 0.473 |
| Thickness ~1 | -0.040 | 0.036 | -0.108, 0.033 | -1.090 | 0.276 |
| Leaf toughness ~1 | 0.017 | 0.041 | -0.062, 0.099 | 0.414 | 0.679 |
| GDD ~1 | 0.001 | 0 | 0.001, 0.001 | -- | -- |
| Indirect LMA effect | 0.149 | 0.042 | 0.070, 0.234 | 3.588 | < 0.001 |
| Indirect Thickness effect | 0.198 | 0.050 | 0.103, 0.301 | 3.963 | < 0.001 |
| Total effect | 0.522 | 0.064 | 0.048, 0.299 | 2.718 | 0.007 |

**Table S10** Model parameters from mediation analysis of growing degree days on leaf strength separately by genetic subpopulation. 95% CIs were calculated via 5000 bootstrapping iterations. The indirect LMA effect is the product of the LMA ~ GDD effect and the Leaf strength ~ LMA effect; the indirect thickness effect is the product of the Thickness ~ GDD effect and the Leaf strength ~ Thickness effect. The total effect is the sum of the two indirect effects and the direct Leaf strength ~ GDD effect. ~ indicates a hypothesized causal relationship; ~~ indicates a hypothesized non-causal relationship.

| | **Path** | **Estimate** | **Std Error** | **95% CIs** | **z** | ***P*** |
|---|---|---|---|---|---|---|
| Atlantic | LMA ~ GDD | 0.621 | 0.081 | 0.469, 0.785 | 7.681 | < 0.001 |
| | Thickness ~ GDD | 0.506 | 0.072 | 0.370, 0.655 | 7.025 | < 0.001 |
| | Leaf strength ~ LMA | 0.195 | 0.124 | 0.005, 0.489 | 1.576 | 0.115 |
| | Leaf strength ~ Thickness | 0.294 | 0.090 | 0.094, 0.450 | 3.263 | 0.001 |
| | Leaf strength ~ GDD | 0.396 | 0.063 | 0.274, 0.521 | 6.307 | < 0.001 |
| | LMA ~~ Thickness | 0.220 | 0.045 | 0.145, 0.326 | 4.935 | < 0.001 |
| | LMA ~~ LMA | 0.368 | 0.048 | 0.285, 0.479 | 7.673 | < 0.001 |
| | Thickness ~~ Thickness | 0.440 | 0.065 | 0.325, 0.590 | 6.731 | < 0.001 |
| | Leaf strength ~~ Leaf strength | 0.194 | 0.061 | 0.107, 0.356 | 3.153 | 0.002 |
| | GDD ~~ GDD | 0.426 | 0 | 0.426, 0.426 | -- | -- |
| | LMA ~1 | -0.228 | 0.062 | -0.349, -0.105 | -3.651 | < 0.001 |
| | Thickness ~1 | -0.253 | 0.066 | -0.376, -0.120 | -3.834 | < 0.001 |
| | Leaf strength ~1 | -0.077 | 0.065 | -0.195, 0.064 | -1.182 | 0.237 |
| | GDD ~1 | -0.432 | 0 | -0.432, -0.432 | -- | -- |
| | Indirect LMA effect | 0.121 | 0.080 | 0.003, 0.327 | 1.510 | 0.131 |
| | Indirect Thickness effect | 0.149 | 0.049 | 0.053, 0.245 | 3.033 | 0.002 |
| | Total effect | 0.666 | 0.066 | 0.552, 0.811 | 10.113 | < 0.001 |
| Gulf | LMA ~ GDD | -0.031 | 0.114 | -0.261, 0.181 | -0.276 | 0.783 |
| | Thickness ~ GDD | 0.251 | 0.106 | 0.039, 0.457 | 2.368 | 0.018 |
| | Leaf strength ~ LMA | 0.169 | 0.068 | 0.033, 0.298 | 2.492 | 0.013 |
| | Leaf strength ~ Thickness | 0.336 | 0.085 | 0.175, 0.510 | 3.948 | < 0.001 |
| | Leaf strength ~ GDD | 0.069 | 0.095 | -0.120, 0.255 | 0.724 | 0.469 |
| | LMA ~~ Thickness | 0.254 | 0.050 | 0.163, 0.362 | 5.028 | < 0.001 |
| | LMA ~~ LMA | 0.720 | 0.092 | 0.567, 0.941 | 7.854 | < 0.001 |
| | Thickness ~~ Thickness | 0.692 | 0.089 | 0.534, 0.886 | 7.790 | < 0.001 |
| | Leaf strength ~~ Leaf strength | 0.606 | 0.071 | 0.486, 0.774 | 8.487 | < 0.001 |
| | GDD ~~ GDD | 0.348 | 0 | 0.348, 0.348 | -- | -- |
| | LMA ~1 | 0.773 | 0.119 | 0.554, 1.019 | 6.517 | < 0.001 |
| | Thickness ~1 | 0.486 | 0.104 | 0.291, 0.701 | 4.680 | < 0.001 |
| | Leaf strength ~1 | 0.403 | 0.101 | 0.207, 0.602 | 4.014 | < 0.001 |
| | GDD ~1 | 0.794 | 0 | 0.794, 0.794 | -- | -- |
| | Indirect LMA effect | -0.005 | 0.021 | -0.058, 0.031 | -0.251 | 0.802 |
| | Indirect Thickness effect | 0.084 | 0.044 | 0.015, 0.189 | 1.908 | 0.056 |
| | Total effect | 0.148 | 0.094 | -0.037, 0.328 | 1.578 | 0.115 |

| | | | | | | |
|---|---|---|---|---|---|---|
| Midwest | LMA ~ GDD | 0.495 | 0.143 | 0.226, 0.787 | 3.454 | < 0.001 |
| | Thickness ~ GDD | 0.377 | 0.160 | 0.057, 0.689 | 2.353 | 0.019 |
| | Leaf strength ~ LMA | 0.231 | 0.070 | 0.091, 0.369 | 3.271 | 0.001 |
| | Leaf strength ~ Thickness | 0.287 | 0.104 | 0.098, 0.507 | 2.755 | 0.006 |
| | Leaf strength ~ GDD | -0.015 | 0.115 | -0.247, 0.205 | -0.130 | 0.896 |
| | LMA ~~ Thickness | 0.158 | 0.041 | 0.094, 0.260 | 3.896 | < 0.001 |
| | LMA ~~ LMA | 0.342 | 0.066 | 0.233, 0.494 | 5.198 | < 0.001 |
| | Thickness ~~ Thickness | 0.290 | 0.053 | 0.204, 0.423 | 5.426 | < 0.001 |
| | Leaf strength ~~ Leaf strength | 0.098 | 0.018 | 0.072, 0.148 | 5.506 | < 0.001 |
| | GDD ~~ GDD | 0.121 | 0 | 0.121, 0.121 | -- | -- |
| | LMA ~1 | -0.332 | 0.114 | -0.562, -0.113 | -2.898 | 0.004 |
| | Thickness ~1 | -0.404 | 0.148 | -0.702, -0.120 | -2.723 | 0.006 |
| | Leaf strength ~1 | -0.491 | 0.089 | -0.648, -0.297 | -5.521 | < 0.001 |
| | GDD ~1 | -0.807 | 0 | -0.807, -0.807 | -- | -- |
| | Indirect LMA effect | 0.114 | 0.046 | 0.045, 0.234 | 2.492 | 0.013 |
| | Indirect Thickness effect | 0.108 | 0.057 | 0.027, 0.264 | 1.904 | 0.057 |
| | Total effect | 0.208 | 0.112 | -0.014, 0.427 | 1.859 | 0.063 |

**Table S11** Model parameters from mediation analysis of growing degree days on leaf toughness separately by genetic subpopulation. 95% CIs were calculated via 5000 bootstrapping iterations. The indirect LMA effect is the product of the LMA ~ GDD effect and the Leaf toughness ~ LMA effect; the indirect thickness effect is the product of the Thickness ~ GDD effect and the Leaf toughness ~ Thickness effect. The total effect is the sum of the two indirect effects and the direct Leaf toughness ~ GDD effect. ~ indicates a hypothesized causal relationship; ~~ indicates a hypothesized non-causal relationship.

| | **Path** | **Estimate** | **Std Error** | **95% CIs** | **z** | ***P*** |
|---|---|---|---|---|---|---|
| Atlantic | LMA ~ GDD | 0.621 | 0.081 | 0.470, 0.789 | 7.674 | < 0.001 |
| | Thickness ~ GDD | 0.506 | 0.072 | 0.371, 0.657 | 7.012 | < 0.001 |
| | Leaf toughness ~ LMA | 0.136 | 0.126 | -0.107, 0.389 | 1.083 | 0.279 |
| | Leaf toughness ~ Thickness | 0.181 | 0.122 | -0.063, 0.412 | 1.485 | 0.138 |
| | Leaf toughness ~ GDD | 0.193 | 0.110 | -0.025, 0.406 | 1.750 | 0.080 |
| | LMA ~~ Thickness | 0.220 | 0.044 | 0.147, 0.326 | 4.978 | < 0.001 |
| | LMA ~~ LMA | 0.368 | 0.048 | 0.284, 0.476 | 7.677 | < 0.001 |
| | Thickness ~~ Thickness | 0.440 | 0.065 | 0.327, 0.586 | 6.759 | < 0.001 |
| | Leaf toughness ~~ Leaf toughness | 0.506 | 0.062 | 0.400, 0.643 | 8.143 | < 0.001 |
| | GDD ~~ GDD | 0.426 | 0 | 0.426, 0.426 | -- | -- |
| | LMA ~1 | -0.228 | 0.063 | -0.348, -0.100 | -3.601 | < 0.001 |
| | Thickness ~1 | -0.253 | 0.066 | -0.374, -0.113 | -3.841 | < 0.001 |
| | Leaf toughness ~1 | -0.190 | 0.074 | -0.327, -0.036 | -2.558 | 0.011 |
| | GDD ~1 | -0.432 | 0 | -0.432, -0.432 | -- | -- |
| | Indirect LMA effect | 0.085 | 0.079 | -0.064, 0.251 | 1.072 | 0.284 |
| | Indirect Thickness effect | 0.091 | 0.063 | -0.030, 0.219 | 1.455 | 0.146 |
| | Total effect | 0.369 | 0.085 | 0.200, 0.532 | 4.357 | < 0.001 |
| Gulf | LMA ~ GDD | -0.031 | 0.111 | -0.253, 0.180 | -0.283 | 0.777 |
| | Thickness ~ GDD | 0.251 | 0.104 | 0.047, 0.462 | 2.404 | 0.016 |
| | Leaf toughness ~ LMA | 0.160 | 0.081 | 0.003, 0.324 | 1.971 | 0.049 |
| | Leaf toughness ~ Thickness | 0.292 | 0.104 | 0.095, 0.503 | 2.809 | 0.005 |
| | Leaf toughness ~ GDD | 0.038 | 0.123 | -0.206, 0.278 | 0.309 | 0.757 |
| | LMA ~~ Thickness | 0.254 | 0.051 | 0.162, 0.361 | 4.962 | < 0.001 |
| | LMA ~~ LMA | 0.720 | 0.089 | 0.569, 0.929 | 8.063 | < 0.001 |
| | Thickness ~~ Thickness | 0.692 | 0.089 | 0.534, 0.896 | 7.733 | < 0.001 |
| | Leaf toughness ~~ Leaf toughness | 0.879 | 0.086 | 0.735, 1.081 | 10.169 | < 0.001 |
| | GDD ~~ GDD | 0.348 | 0 | 0.348, 0.348 | -- | -- |
| | LMA ~1 | 0.773 | 0.117 | 0.557, 1.014 | 6.633 | < 0.001 |
| | Thickness ~1 | 0.486 | 0.104 | 0.286, 0.697 | 4.670 | < 0.001 |
| | Leaf toughness ~1 | 0.243 | 0.134 | -0.011, 0.515 | 1.810 | 0.070 |
| | GDD ~1 | 0.794 | 0 | 0.794, 0.794 | -- | -- |
| | Indirect LMA effect | -0.005 | 0.020 | -0.057, 0.028 | -0.254 | 0.800 |
| | Indirect Thickness effect | 0.073 | 0.042 | 0.014, 0.185 | 1.743 | 0.081 |
| | Total effect | 0.106 | 0.124 | -0.131, 0.356 | 0.859 | 0.390 |

| | | | | | | |
|---|---|---|---|---|---|---|
| Midwest | LMA ~ GDD | 0.495 | 0.143 | 0.217, 0.775 | 3.461 | < 0.001 |
| | Thickness ~ GDD | 0.377 | 0.160 | 0.068, 0.700 | 2.363 | 0.018 |
| | Leaf toughness ~ LMA | 0.268 | 0.152 | -0.004, 0.599 | 1.771 | 0.077 |
| | Leaf toughness ~ Thickness | 0.377 | 0.177 | 0.044, 0.736 | 2.131 | 0.033 |
| | Leaf toughness ~ GDD | 0.012 | 0.248 | -0.466, 0.519 | 0.048 | 0.962 |
| | LMA ~~ Thickness | 0.158 | 0.040 | 0.092, 0.254 | 3.901 | < 0.001 |
| | LMA ~~ LMA | 0.342 | 0.066 | 0.233, 0.498 | 5.151 | < 0.001 |
| | Thickness ~~ Thickness | 0.290 | 0.054 | 0.201, 0.420 | 5.395 | < 0.001 |
| | Leaf toughness ~~ Leaf toughness | 0.535 | 0.091 | 0.388, 0.761 | 5.880 | < 0.001 |
| | GDD ~~ GDD | 0.121 | 0 | 0.121, 0.121 | -- | -- |
| | LMA ~1 | -0.332 | 0.114 | -0.568, -0.120 | -2.921 | 0.003 |
| | Thickness ~1 | -0.404 | 0.146 | -0.691, -0.120 | -2.770 | 0.006 |
| | Leaf toughness ~1 | 0.060 | 0.193 | -0.304, 0.462 | 0.313 | 0.754 |
| | GDD ~1 | -0.807 | 0 | -0.807, -0.807 | -- | -- |
| | Indirect LMA effect | 0.133 | 0.092 | 0.008, 0.385 | 1.440 | 0.150 |
| | Indirect Thickness effect | 0.142 | 0.083 | 0.025, 0.387 | 1.710 | 0.087 |
| | Total effect | 0.287 | 0.244 | -0.170, 0.785 | 1.179 | 0.238 |

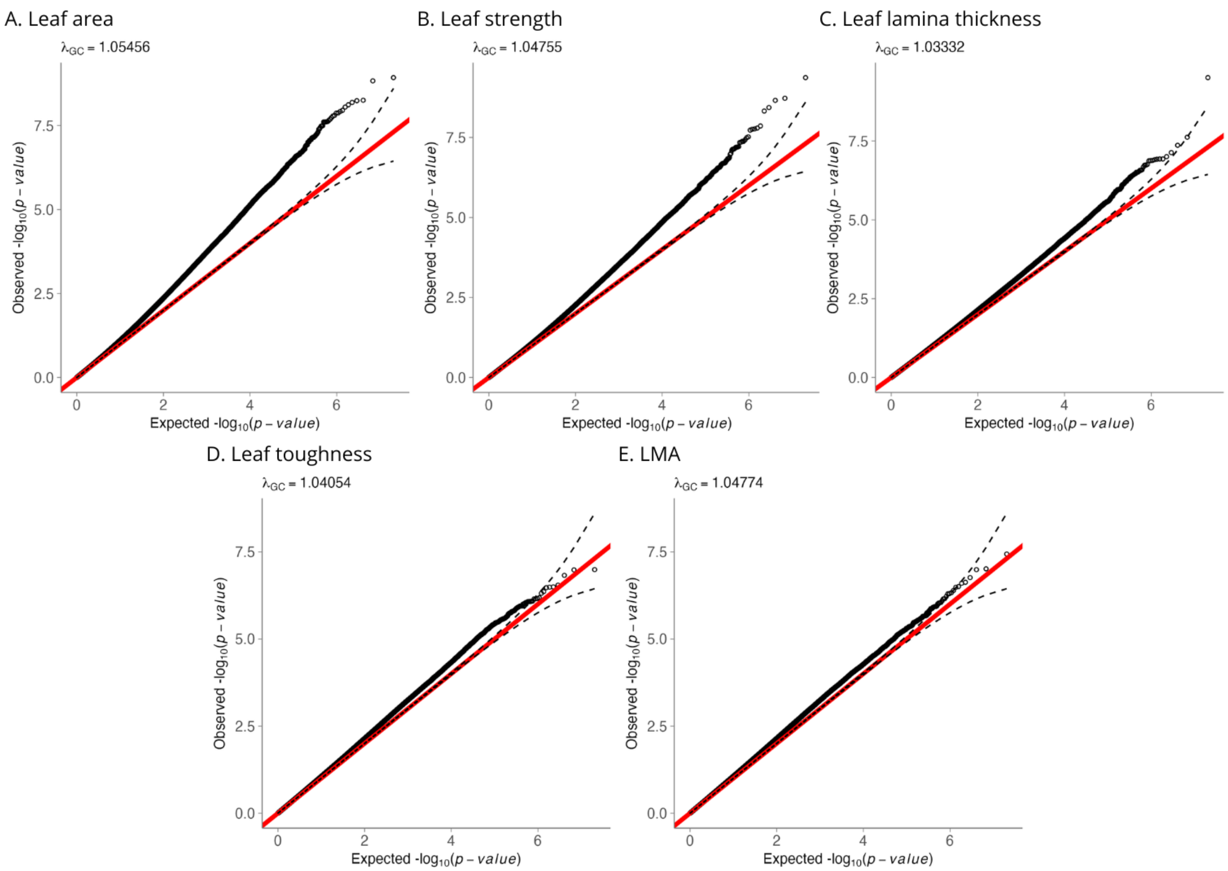


**Fig. S1** Observed vs. expected p-values for single-trait GWA. The red line represents a 1:1 ratio.

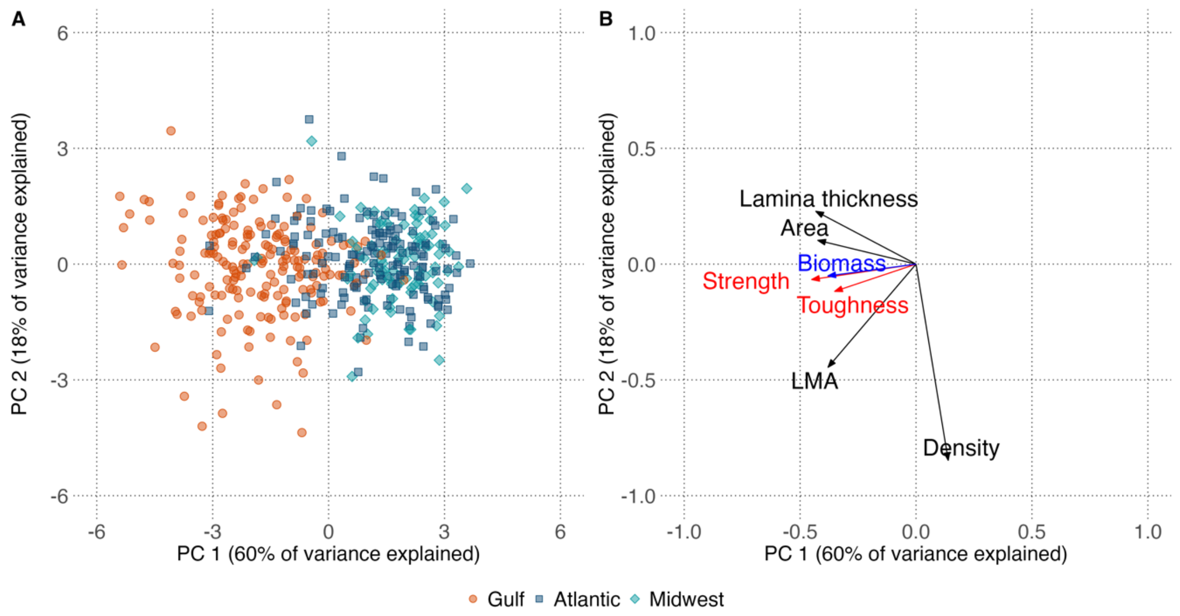


**Fig. S2** PCA of phenotypes. A) shows each individual in the space of PC1 and PC2 with color and shape representing the three different subpopulations. B shows the loading of each phenotype onto PC1 and PC2, with black arrows representing leaf morphological traits, red arrows representing leaf tensile resistance traits, and a blue arrow representing biomass (fitness).

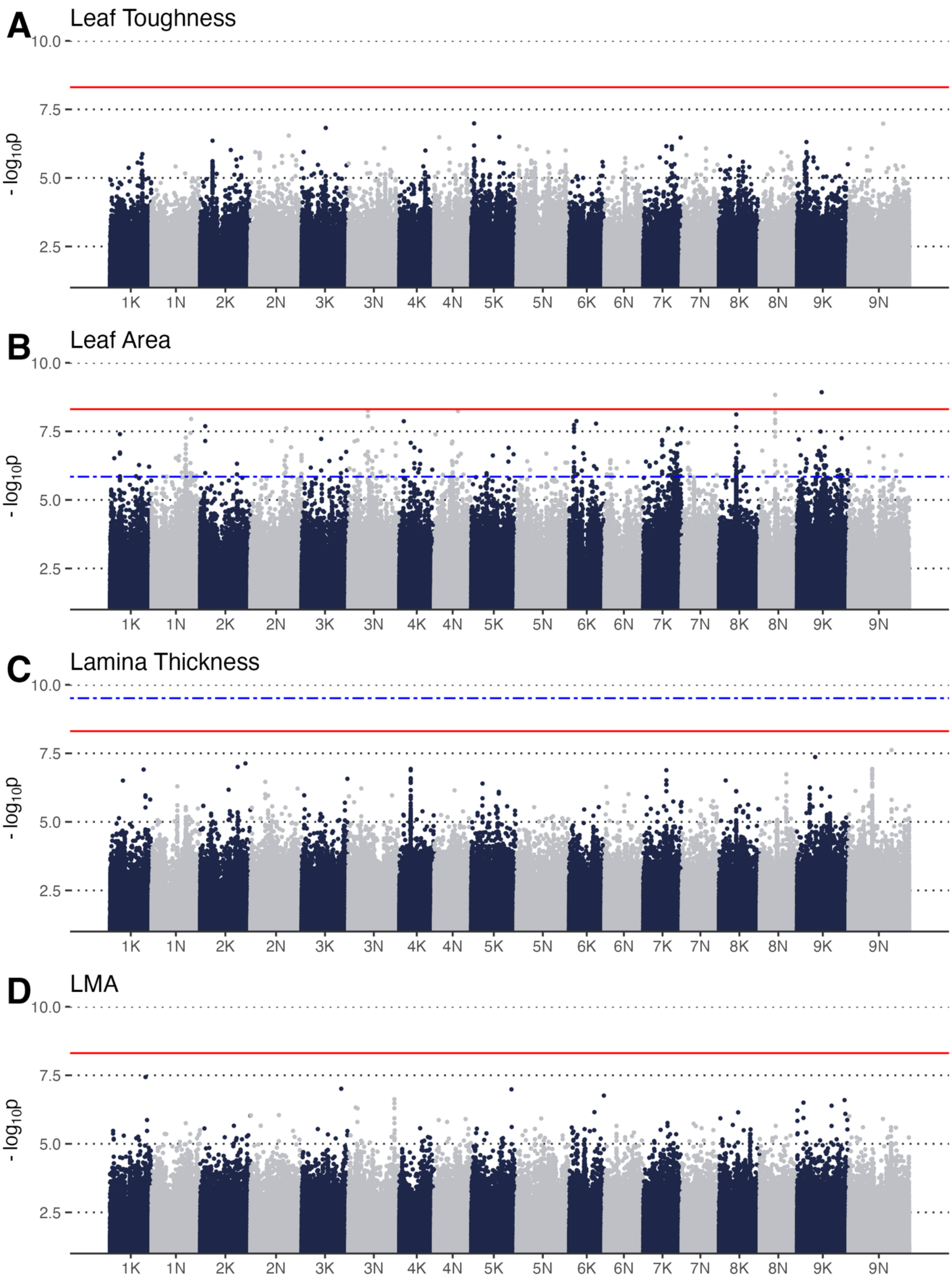
A
Leaf Toughness
B
Leaf Area
C
Lamina Thickness
D
LMA
10.0
7.5
5.0
2.5
- log10p
1K 1N 2K 2N 3K 3N 4K 4N 5K 5N 6K 6N 7K 7N 8K 8N 9K 9N

**Fig. S3** Manhattan plots showing the associations between single nucleotide polymorphisms (SNPs) and leaf traits. The solid red lines represent a genome-wide Bonferroni-adjusted significance threshold and the dashed blue lines below represent a genome-wide 5% false discovery rate threshold; SNPs above these lines have a significant statistical association with leaf traits ($-\log_{10}$ *P*-value). Manhattan plots without a dashed blue line represent a GWAS that did not have any significant SNPs associated with the respective trait. For plotting purposes, we removed all SNPs whose $-\log_{10}$ p-value $< 1$.

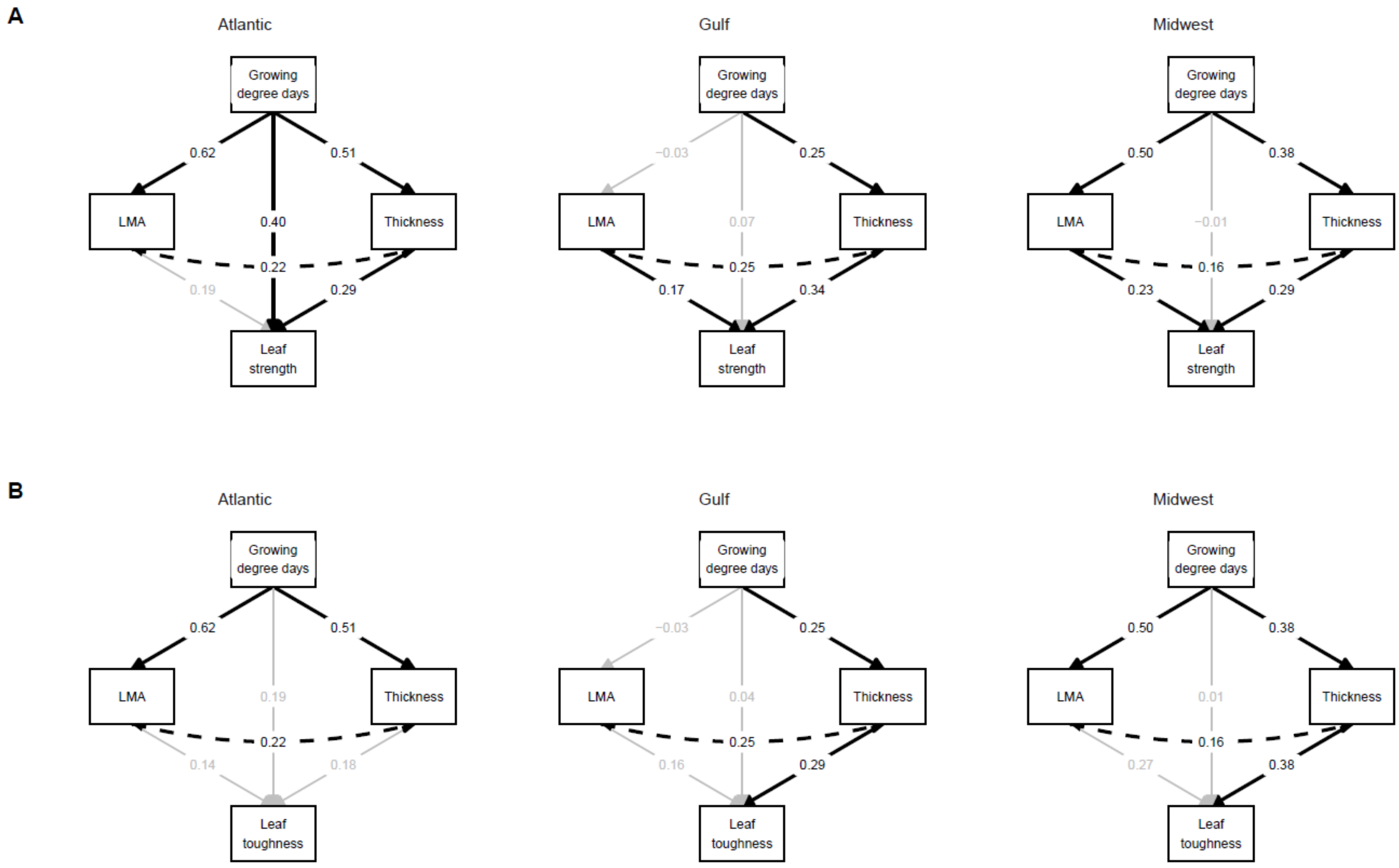
A
Atlantic
Growing degree days
LMA
Thickness
Leaf strength
0.62
0.51
0.40
0.22
0.19
0.29
Gulf
Growing degree days
LMA
Thickness
Leaf strength
−0.03
0.25
0.07
0.25
0.17
0.34
Midwest
Growing degree days
LMA
Thickness
Leaf strength
0.50
0.38
−0.01
0.16
0.23
0.29
B
Atlantic
Growing degree days
LMA
Thickness
Leaf toughness
0.62
0.51
0.19
0.22
0.14
0.18
Gulf
Growing degree days
LMA
Thickness
Leaf toughness
−0.03
0.25
0.04
0.25
0.16
0.29
Midwest
Growing degree days
LMA
Thickness
Leaf toughness
0.50
0.38
0.01
0.16
0.27
0.38

**Fig. S4** Direct and indirect effects of growing degree days, a proxy for growing season length, on A) leaf strength and B) leaf toughness, separately by genetic subpopulation, showing the importance of two leaf morphological traits, LMA and leaf thickness. All coefficients are standardized. Black lines denote significant relationships ($p < 0.05$); grey lines denote non-significant relationships. Growing degree days was mean-centered. All other variables were standardized such that across all plants, mean = 0 and sd = 1.

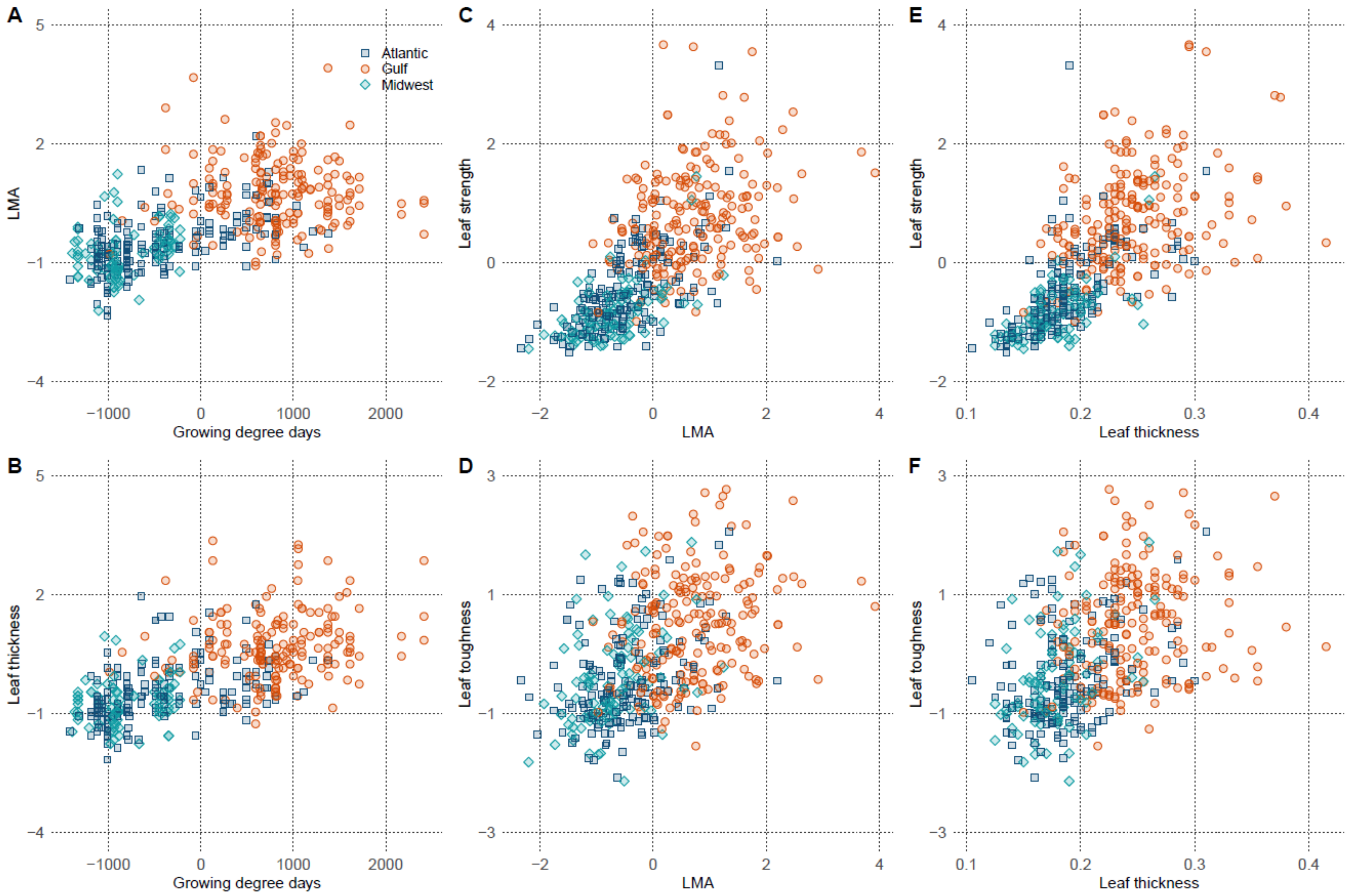

A
B
C
D
E
F
Atlantic
Gulf
Midwest
LMA
Leaf thickness
Leaf strength
Leaf toughness
Growing degree days
LMA
Leaf thickness
5
2
−1
−4
4
3
0
−2
1
−3
−1000
0
1000
2000
−2
0
2
4
0.1
0.2
0.3
0.4

**Fig. S5** Direct and indirect effects of growing degree days, a proxy for growing season length, on leaf strength and leaf toughness, separately by genetic subpopulation and mediated by two leaf morphological traits, LMA and leaf thickness. All coefficients are standardized. Growing degree days was mean-centered. All other variables were standardized such that across all plants, mean = 0 and sd = 1.
